# Novel gene-specific translation mechanism of dysregulated, chronic inflammation reveals promising, multifaceted COVID-19 therapeutics

**DOI:** 10.1101/2020.11.14.382416

**Authors:** Li Wang, Adil Muneer, Ling Xie, Feng Zhang, Bing Wu, Liu Mei, Erik M Lenarcic, Emerald Hillary Feng, Juan Song, Yan Xiong, Xufen Yu, Charles Wang, Ciprian Gheorghe, Karina Torralba, Jeanette Gowen Cook, Yisong Y. Wan, Nathaniel John Moorman, Hongjun Song, Jian Jin, Xian Chen

## Abstract

Hyperinflammation and lymphopenia provoked by SARS-CoV-2-activated macrophages contribute to the high mortality of Coronavirus Disease 2019 (COVID-19) patients. Thus, defining host pathways aberrantly activated in patient macrophages is critical for developing effective therapeutics. We discovered that G9a, a histone methyltransferase that is overexpressed in COVID-19 patients with high viral load, activates translation of specific genes that induce hyperinflammation and impairment of T cell function or lymphopenia. This noncanonical, pro-translation activity of G9a contrasts with its canonical epigenetic function. In endotoxin-tolerant (ET) macrophages that mimic conditions which render patients with pre-existing chronic inflammatory diseases vulnerable to severe symptoms, our chemoproteomic approach with a biotinylated inhibitor of G9a identified multiple G9a-associated translation regulatory pathways that were upregulated by SARS-CoV-2 infection. Further, quantitative translatome analysis of ET macrophages treated progressively with the G9a inhibitor profiled G9a-translated proteins that unite the networks associated with viral replication and the SARS-CoV-2-induced host response in severe patients. Accordingly, inhibition of G9a-associated pathways produced multifaceted, systematic effects, namely, restoration of T cell function, mitigation of hyperinflammation, and suppression of viral replication. Importantly, as a host-directed mechanism, this G9a-targeted, combined therapeutics is refractory to emerging antiviral-resistant mutants of SARS-CoV-2, or any virus, that hijacks host responses.

## Introduction

The Coronavirus Disease (COVID-19) pandemic, caused by severe acute respiratory syndrome coronavirus-2 (SARS-CoV-2), is an unprecedented global public health crisis. As of October 16, 2020, the virus has claimed over 1.1-million lives. High mortality is observed for SARS-CoV-2-infected patients who have preexisting chronic conditions such as recovery from sepsis, chronic pulmonary diseases, metabolic diseases (e.g., diabetes), asthma, cardiovascular diseases, thrombosis, chronic liver disease and cirrhosis, and cancer^1,2^. A dysregulated immune system or hyperinflammatory response to SARS-CoV-2 infection is the leading cause of severe illness and mortality.^3,4^ Correspondingly, excessive release of inflammatory factors in a ‘cytokine storm’ aggravates acute respiratory distress syndrome (ARDS) or widespread tissue damage, which results in respiratory or multi-organ failure and death. The immunopathologic characteristics of COVID-19 include reduction and functional exhaustion of T cells (lymphopenia) and increased levels of serum cytokines (hyperinflammation)^5^. Ordinarily, macrophages regulate the innate immune response to viral threats by producing inflammatory molecules that activate T cells for viral containment and clearance. However, macrophage function/activation is dysregulated in severe COVID-19 patients^6,7^, and proinflammatory monocyte-derived macrophages are abundant in their bronchoalveolar lavage fluids.^8^ These observations indicated the crucial effects of dysregulated macrophages in SARS-CoV-2 immunopathogenesis and associated cytokine storms. Therapeutic options have been severely restrained by a lack of understanding of the pathways and mechanisms that trigger the SARS-CoV-2-induced hyperinflammatory response.

As the systemic cytokine profiles in severe COVID-19 patients showed similarities to profiles in macrophage activation syndromes^9^, particularly viral sepsis^10^, we investigated the mechanistic causes of SARS-CoV-2-induced immunopathology in macrophages. We used macrophage cells that had acquired endotoxin (lipopolysaccharide) tolerance (ET). ET is the common immunopathological background of COVID-19 vulnerable groups that have pre-existing chronic inflammatory diseases^11,12^. Genome-wide CRISPR screening revealed the crucial role of epigenetic regulation involving chromatin remodeling and histone modification in SARS-CoV-2 infection^13^. Correspondingly, the histone methyltransferases G9a and G9a-like protein (GLP; hereafter G9a will represent both proteins) that are constitutively activated in ET^14^ showed upregulated expression in COVID-19 patients with high viral load^15^. Further, we showed that inhibition of G9a enzyme activity mitigated or reversed ET,^14^ which implicated active G9a in ET-related, SARS-CoV-2-induced pathogenesis. Thus, to dissect G9a-associated pathways and mechanisms in ET, we used our chromatin activity-based chemoproteomic (ChaC) approach, which consists of a biotinylated G9a inhibitor UNC0965^16^, to capture ET-phenotypic G9a protein complexes and identify their constituents by mass spectrometry (MS). Notably, ChaC-MS is superior to conventional immunoprecipitation-MS^17,18^ that captures protein complexes based only on epitope abundance. ChaC-MS identified protein partners of constitutively active G9a that together represent the ET phenotype. The ET macrophage partners of constitutively active G9a included many proteins associated primarily with translation initiation and elongation, RNA modification and processing, and ribosome biogenesis. Strikingly, these G9a interactors involving translational regulation were also identified in cellular pathways that are upregulated or ‘reshaped’ by SARS-CoV-2^19^. Notably, most cofactors of the (m^6^A) RNA methylase METTL3^20^ were among these ET-phenotypic G9a interactors. METTL3 appears to promote translation of specific oncogenes,^20^ and is implicated in inflammation.^21–23^ Thus, we hypothesized that G9a exerts a noncanonical (nonepigenetic silencing) function, in conjunction with METTL3 and *other* translation regulators to promote translation of mRNAs that establish the ET phenotype. As the first step to characterize this novel function of G9a, we used m^6^A RNA immunoprecipitation and label-free quantitative proteomics^24–26^ to identify subsets of m^6^A-tagged mRNAs whose translation showed a dual dependence on G9a and METTL3 in ET macrophages. Among these subsets, the sepsis-characteristic overexpression of an immune checkpoint protein CD274 (programmed cell death-ligand 1 or PD-L1)^27^ and multiple chemokine receptors^28^ were co-upregulated by G9a and METTL3. Crucially, depletion of G9a and METTL3 restored T cell function and reduced the survival of macrophages. Further, we determined proteome-wide translatome changes following pharmaceutical inhibition or genetic knock-out of G9a. This analysis revealed that, from a broader view extended from the G9a-METTL3 translation regulatory axis, G9a activated the translation of specific proteins that were identified in diverse SARS-CoV-2 upregulated pathways^19^ or in the PBMCs^29^ or sera of severe COVID patients^6^. Accordingly, treatment of ET macrophages with a G9a inhibitor mitigated the elevated expression of COVID-19-characteristic proteins. Additionally, because the histone methyltransferase Ezh2 was identified as an ET-specific interactor of G9a, we treated ET macrophages with an Ezh2 inhibitor and similarly observed mitigation of elevated COVID-19-characteristic proteins.

From a systems view, we discovered the G9a-associated mechanism of SARS-CoV-2 immunopathogenesis in which constitutively active G9a promotes the translation of genes that comprise diverse pathways involved in host-virus interactions or viral replication^30^ and the host response to SARS-CoV-2 infection.^19^ Thus, G9a-target therapy for COVID-19 has potential multifaceted effects to inhibit host hyperinflammation, restore T cell function, and inhibit virus replication. Thus, proteins showing G9a- or Ezh2-dependent overexpression could serve as biomarkers for stratification of COVID-19 patients responsive to G9a- or Ezh2-target therapy. Our discovery of the mechanism by which the G9a/Ezh2 inhibitors reverse SARS-CoV-2 dysregulated inflammation will facilitate design of effective, precision therapies that may improve or facilitate patient survival or recovery from COVID-19 and other chronic inflammatory diseases.

## RESULTS

### Constitutively active G9a is implicated in SARS-CoV-2 upregulated translation pathways

We found that mRNAs which encode major components of the G9a/GLP (EHMT2/EHMT1)-associating complex were overexpressed in COVID-19 patients with increasing SARS-CoV-2 load (**Extended Data Fig. 1a**)^15^. Accordingly, we found by top-down mass spectrometry (MS), ChIP-PCR, and ChaC chemoprobe pull-down that the methylation activity of G9a was constitutively higher in chronically inflamed or endotoxin-tolerant (TL or ET) macrophages compared with acutely inflamed (NL) cells.^14^ Thus, we performed label-free quantitation (LFQ),^25,26^ UNC0965 ChaC experiments^16^ on mouse Raw 264.7 macrophages under nonstimulated (N) and different inflammatory conditions (NL, TL) (**Fig. 1a**). On the basis of LFQ ratios that are proportional to the relative binding of individual proteins to G9a in TL versus NL/N, we identified >1000 proteins that showed consistently enhanced interaction with G9a in TL/ET macrophages (**Extended Data Table 1**).

**Fig.1.**
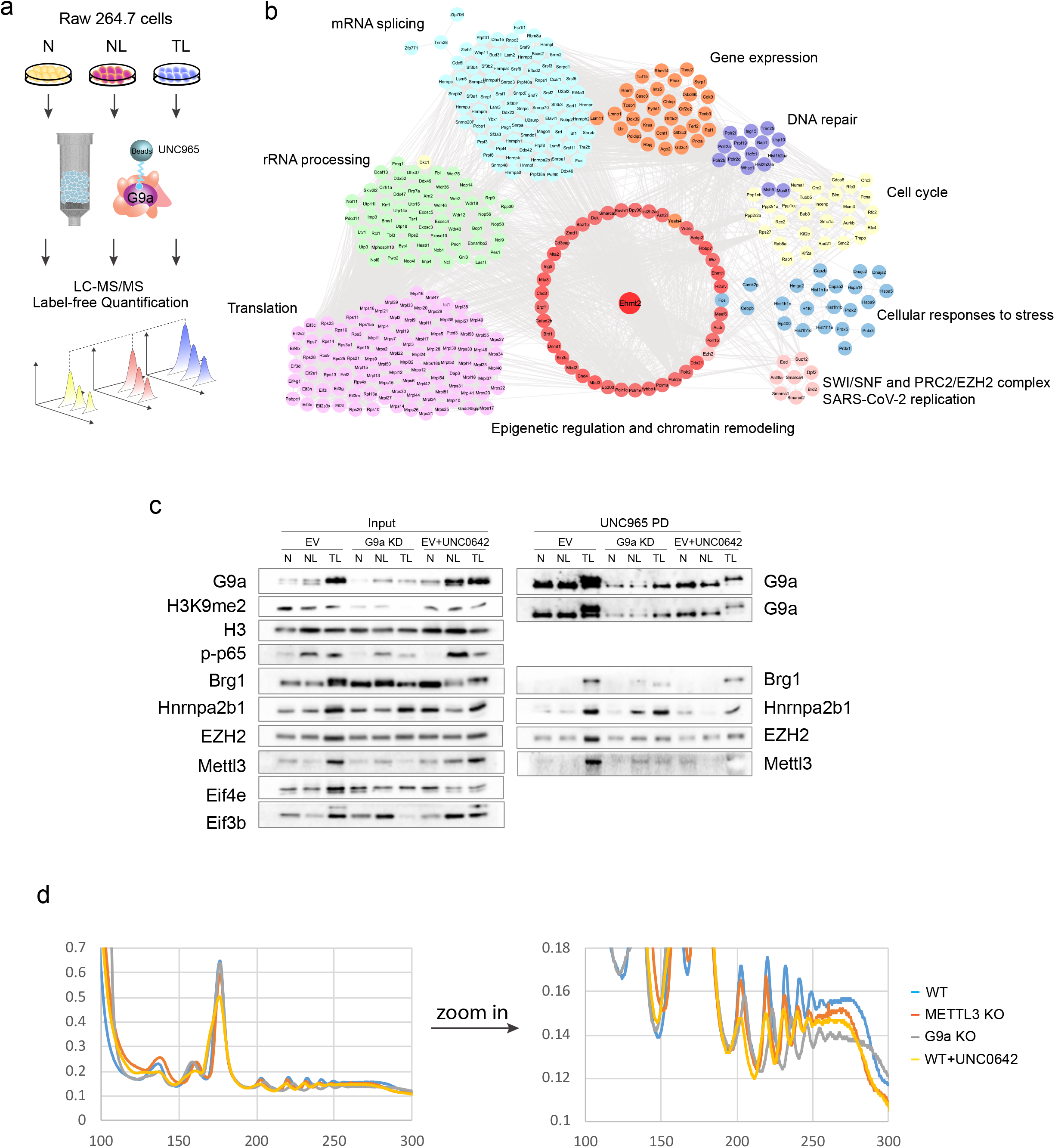
Constitutively active G9a is implicated in SARS-CoV-2 upregulated translation pathways in ET. **a**. Schematic of LFQ ChaC-MS dissection of G9a interactome in ET macrophages. **b**. Over-represented functional pathways and networks of G9a interactors in chronically inflamed macrophages (TL/ET). Physical interactions were curated from StringApp, Reactome in Cytoscape (v3.8.1). **c**. Immunoblot analysis of ET-specific associations with endogenous G9a for METTL3 and major translation regulators. The Raw264.7 macrophage lines either stably expressing shRNA for G9a knock-down (KD), or empty vector (EV) for G9a wild type. ‘EV+UNC0642’ refers to G9a inhibitor (UNC0642) treatment. (left) inputs, and (right) UNC0965 pull-down (PD). Some interactors were pulled down from the G9a KD cells because of residual G9a. **d**. Polysome analysis of G9a- or METTL3-dependent protein synthesis. Absorbance profile of sucrose density gradients showing the location of 40S and 60S ribosomal subunits, 80S monosomes, and polysomes. Wild-type (blue line), G9a knock-out or ko (gray line), and METTL3 ko (orange line), Wild-type with UNC0642 treatment(yellow line).

We used STRING (Search Tool for the Retrieval of Interacting Genes) to identify enriched functional pathways/processes that were represented by these ET-specific G9a interactors (**Fig. 1b**). First, in agreement with G9a’s canonical epigenetic regulatory function, we identified proteins associated with chromatin remodeling and histone modification. These chromatin proteins included the SWI/SNF remodeling complex and BRD4, which were characterized by siRNA and genome-wide CRISPR screens as essential host factors for SARS-specific and pan-coronavirus infection, respectively^13,30^ (**Fig. 1b**). These results implicated the ET-phenotypic G9a interactome in SARS-CoV-2 immunopathogenesis. Surprisingly, ChaC data from ET macrophages predicted that, via ET-specific interactions with some representative proteins, G9a may have a noncanonical (nonepigenetic) function in major translation regulatory processes. Strikingly, fifty-three G9a interactors that we identified were found by Bojkova et al. in SARS-CoV-2 upregulated pathways associated with translation initiation/elongation, alterative splicing, RNA processing, nucleic acid metabolism, and ribosome biogenesis^19^ (**Extended Data Fig. 1b**). This coincident identification validated the similarity between ET-immunophenotypic G9a pathways and pathways activated by SARS-CoV-2. For example, we identified the splicing factor SF3B1 and the 40S ribosomal protein Rps14 as ET-specific G9a interactors, and Bojkova et al. found that emetine inhibition of Rps14 or pladienolide inhibition of SF3B1 significantly reduced SARS-CoV-2 replication^19^. Like our finding for G9a interactors in cancer patients with poor prognosis^16^, the ChaC-MS results for ET macrophages showed that certain genes upregulated by SARS-CoV-2 infection are fully translated to their encoded proteins as ET-specific G9a interactors. Thus, constitutively active G9a may coordinate these SARS-CoV-2 activated translation pathways.

### Interaction with METTL3 and other translation proteins implicates G9a in translational regulation of chronic inflammation

We noticed that UNC0965 captured most known cofactors of METTL3. METTL3 enhances mRNA translation by interaction with the translation initiation machinery^20^ such as NCBP1/2 (CBP80) and RBM15, HNRNPA2B1, eIF3, and eIF4E. Although METTL3 itself was not identified by MS, our identification of METTL3 cofactors as TL-specific G9a interactors indicated that G9a may regulate translation via interaction with METTL3. We also identified most of the other translation regulatory proteins in Flag-tagged METTL3 pull-down from TL macrophages (**Extended Data Table 2**). For example, ribosomal proteins that bind to actively translated mRNA were identified, including forty-six 39S ribosomal proteins (Mrpl1-57), 28S (Mrps), 40S ribosomal proteins (Rps), 60S ribosomal proteins (RPl), 60S ribosome subunit biogenesis protein NIP7 homolog, ribosome-binding protein 1 (Rrbp1), ribosomal RNA processing proteins (Rrp1, Rrp36, Rrp7a, Rrp8) as well as multiple RNA-binding proteins (**Fig. 1b**). These results from unbiased interactome screening^31^ indicated that G9a and METTL3 function in the same translation regulatory pathways. Immunoblotting showed TL-specific association with endogenous G9a with METTL3 and major translation regulators (**Fig. 1c**), in agreement with the ChaC result. In macrophages with G9a knock-down, UNC0965 pulled down only proteins that interacted with the residual G9a. These results confirmed the new function of G9a in METTL3-mediated translation in ET macrophages.

We then determined by polysome analysis^32^ the effect of G9a and METTL3 on protein synthesis. First, we used CRISPR/Cas9 to knock out (ko) G9a (EHMT2) or METTL3 in macrophages subjected to LPS stimulation to generate cells with different inflammatory conditions (NL, T, or TL). G9a depletion caused reduced protein expression of METTL3 in TL/ET. Also, as indicated by reduced levels of the proinflammatory markers such as phosphor-p65 and phospho-IκB in the macrophages under prolonged LPS stimulation (**Extended Data Fig. 1c**), depletion of either METTL3 or G9a similarly caused reduced ET and resensitized the ET macrophages to LPS stimulation. These results suggested that ET is coregulated by G9a and METTL3. Next, we measured global protein synthesis from polysome abundance in wild type versus G9a ko or METTL3 ko macrophages.^32^ **Fig. 1d** shows that the polysome abundance decreased in G9a ko or G9a inhibitor(UNC0642) treated or METTL3 ko cells compared with abundance in wild type. Thus, depletion of either G9a or METTL3 suppressed protein synthesis that was dependent on G9a and METTL3 in the endotoxin-tolerant macrophages.

### G9a and METTL3 co-upregulate translation of m^6^A-marked mRNA subsets

Because global protein synthesis in ET macrophages required G9a and METTL3, we next sought to identify specific mRNAs whose translation was dependent on G9a and METTL3. Thus, we performed RNA-seq and m^6^A RNA immunoprecipitation-sequencing (MeRIP-Seq) on wild type, G9a ko, and METTL3 ko THP1 macrophages in N, NL, TL. First, in the G9a-depleted macrophages under TL, 440 immune or inflammatory response genes exhibited increased mRNA abundance (**Extended Data Fig. 2a**). This finding of the G9a-suppressed genes aligned with our previous report^14^ that, via interactions with transcriptional repressors such as cMyc, constitutively active G9a suppresses the transcription of proinflammatory genes. Conversely, we identified, in both G9a ko and METTL3 ko cells under TL, 136 genes with decreased mRNA expression compared to wild type cells; these genes are associated mostly with regulation of cell cycle progression, cell proliferation, and antiviral or anti-inflammatory responses. Like the co-existence of G9a and METTL3 in TL-specific translation regulatory complexes, this new finding of G9a/METTL3-co-upregulated genes indicated that G9a and METTL3 cooperate to posttranscriptionally activate expression of pro-survival and anti-inflammatory genes in TL. Further, 62 of 136 G9a/METTL3-co-upregulated genes were tagged with m^6^A (**Fig. 2a**). The Integrative Genomics Viewer plots showed that the m^6^A level of most mRNAs that encode these G9a/METTL3-co-upregulated genes decreased in either G9a ko or METTL3 ko macrophages with TL (**Extended Data Fig. 2b**). Notably, the ‘stabilized’ or ‘actively translated’ mRNA (total input) that was proportional to the amount of m^6^A-tagged transcripts also showed a dependence on G9a and METTL3. We validated by quantitative PCR the G9a- and METTL3-dependence of the abundance of m^6^A mRNA coding these genes (**Extended Data Fig. 2c**).

**Fig.2.**
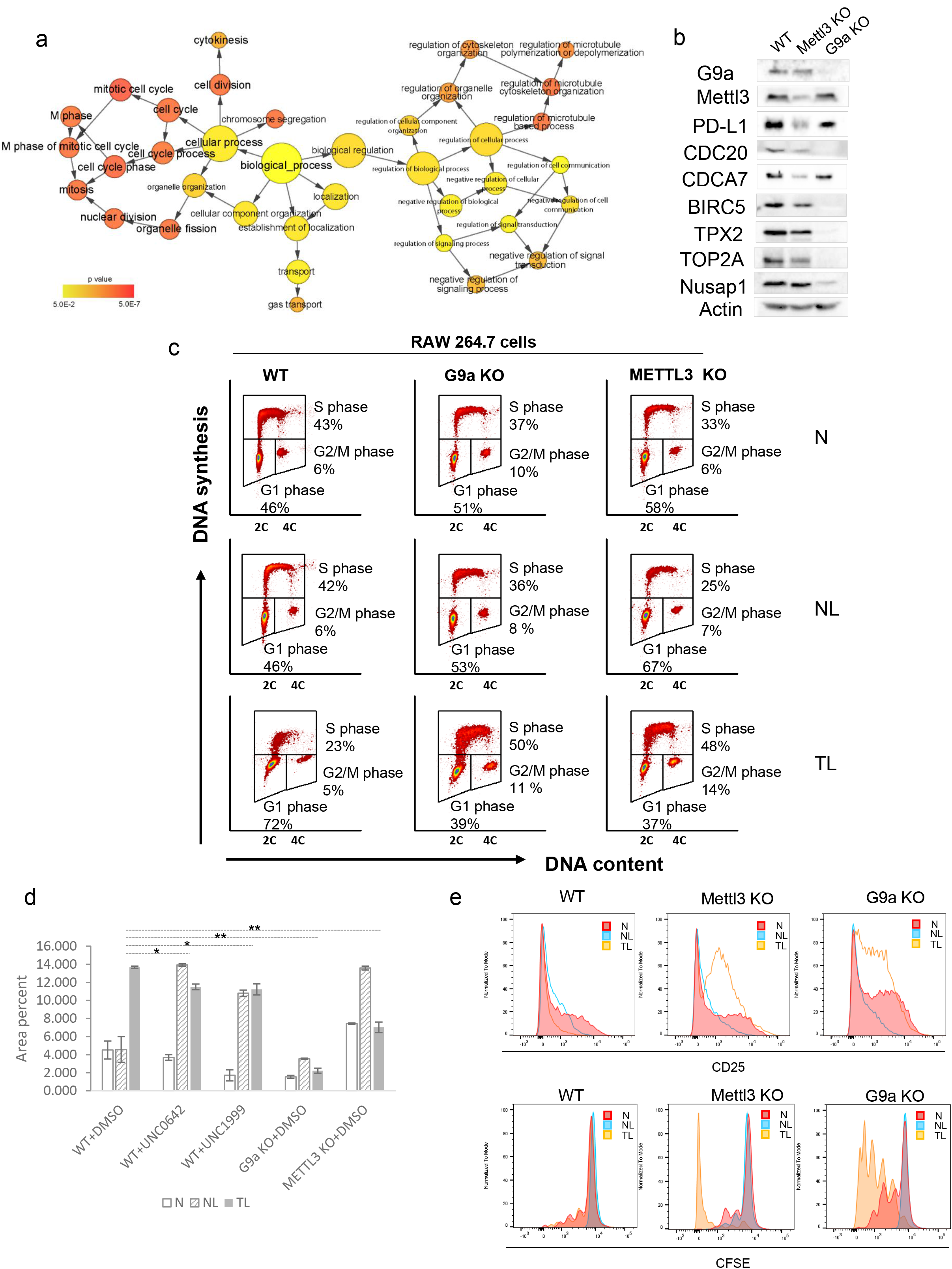
G9a and METTL3 cooperate to dysregulate cell cycle and impair T cell function under ET. **a**. Enriched functional pathways overrepresented by G9a/METTL3-co-downregulated m^6^A mRNA in ET. The network was constructed by BinGO app in Cytoscape. **b**. Immunoblot of protein expression in endotoxin tolerized wild-type versus G9a ko or METTL3 ko THP-1 cells. **c**. LPS-induced cell cycle arrest in endotoxin tolerance condition requires G9A and METTL3. Flow cytometry analysis of the impact of G9a and METTL3 on cell cycle in different inflammatory conditions (*e.g*., N, NL, or TL). Cells were labeled with EdU for 0.5 h before harvesting and analyzed by flow cytometry for DNA content with DAPI and for DNA synthesis with EdU. **d**. Clonogenic survival assay of WT and KO Raw 264.7 cells in different inflammation conditions. WT cells were either treated with DMSO (0.05%) or 1 mM G9a inhibitor (UNC0642) or EZH2 inhibitor (UNC1999). * p<0.05, ** p<0.01. **e**. Depletion of METTL3 or G9a promotes the T cell activation and proliferation under the TL conditions. Histograms obtained from the CD8 T-cell activation (**upper panel**) and proliferation (**lower panel**). The proliferation and activation markers, including CD25 of P14 CD8^+^ T cells, were analyzed by flow cytometry at day 5 and day 6 after coculture with wild type, METTL3 knock-out (ko) or G9a ko RAW 264.7 cells that were untreated (N) or treated with an acute LPS stimulation (NL) and prolonged LPS stimulation (TL), a mimic of ET.

To determine the predominant mechanism responsible for the G9a-mediated translation of specific genes, we examined whether G9a ko affected the m^6^A level of total RNA under N, NL, TL. We observed a relatively reduced m^6^A level in G9a ko and METTL3 ko cells (**Extended Data Fig. 2d**). In addition, we performed LFQ proteomics^26^ to identify proteins that exhibited G9a- or METTL3-dependent expression changes in the same THP macrophage set (e.g., wild type versus G9a ko versus METTL3). Principle component analysis (**Extended Data Fig. 2e**) showed that, in TL, G9a ko or METTL3 ko produced clusters of protein expression profiles that were well separated from the clusters of wild type ET macrophages. This result indicated that depletion of G9a and METTL3 led to characteristic protein expression patterns that represented similar inflammatory phenotypes. Importantly, the G9a/METTL3-co-upregulated m^6^A mRNA or proteins that we identified by nonbiased MeRIP-Seq or LFQ proteomics functionally overlapped with pathways related to antiviral immune response and cell fate determination. Immunoblotting confirmed that the protein expression of select genes in these functional clusters had a dual dependence on G9a and METTL3 (**Fig. 2b**), which indicated that G9a and METTL3 coactivate translation of the same sets of m^6^A-tagged mRNA. In agreement with our ChaC-MS finding of enhanced interactions of G9a with METTL3 and associated translational regulatory proteins (**Fig. 1b**), this epitranscriptomic-to-proteomic correlation for specific genes indicated that G9a is directly involved in gene-specific translation regulatory processes in ET macrophages.

### G9a and METTL3 promote proliferation of ET macrophages that produce organ-damaging inflammatory factors

One cluster of the proteins co-upregulated by G9a and METTL3, BIRC5,^33^ CDC20,^34,35^ NUSAP1,^36^ CDCA7,^37^ TOP2A, and ALCAM, is functionally associated with cell division, mitotic cell cycle, and cell proliferation^36,38–41^ (**Fig. 2a**). This observation not only aligned with the function of METTL3 in promoting cell cycle progression and survival^42^, but, more importantly, the observation implicated constitutively active G9a in promoting METTL3-mediated translation of prosurvival proteins, e.g., METTL3-mediated m^6^A promotes translation of cMyc mRNA and is responsible for cMyc stability in AML cells^43^.

We therefore used flow cytometry to determine the effect of G9a and METTL3 on the cell cycle in different inflammatory conditions. As shown in **Fig. 2c**, conditions that mimicked chronic inflammation (TL/ET) caused a dramatic accumulation of macrophages in G1 compared with nonstimulated (N) or acutely inflamed (NL) conditions (72% vs 46%). This G1 accumulation was accompanied by fewer S phase cells (23% vs 43%). Neither G9a nor METTL3 knockout delayed the G1 phase under the TL condition. Prolonged LPS stimulation (TL) caused these G9a- or METTL3-knockout lines to spend more time in S phase relative to wild type cells and relative to the same cells under acute inflammation conditions. The clonogenic assay (**Fig. 2d**) showed both knockout lines survived less well than controls during TL treatment, and the G1 delay could serve a protective function. Akin to our proteomic finding that G9a/METTL3 coactivate expression of multiple prosurvival proteins, these cell cycle analyses indicated that constitutively active G9a and METTL3 cooperate to restrain cell cycle progression and promote increased survival of the slow-processing macrophages during chronic inflammation, possibly by targeting prosurvival mRNAs for translation specifically in the growth phase (G1).

### G9a and METTL3 co-upregulated ET overexpression of PD-L1, and depletion of G9a or METTL3 restored T cell function

Another cluster of G9a/METTL3-co-upregulated proteins, PD-L1, CX3CR1, and IRF8, are functionally associated with immune checkpoint regulation and antimicrobial response. PD-L1 is overexpressed in sepsis patients with impaired T cell function,^27^ and, likewise, our MeRIP-Seq data showed an increased level of PD-L1 (CD274) m^6^A mRNA under ET **(Extended Data Fig. 2b)**. Although PD-L1 m^6^A mRNA exhibited little dependence on G9a or METTL3 similar to certain METTL3-regulated genes,^20^ **r**esults from LFQ proteomics and immunoblotting consistently showed that the ET overexpression of PD-L1 was dependent on G9a and METTL3 (**Fig. 2b and Extended Data Fig.2e**). Thus, we hypothesized that G9a and METTL3 impair T cell function via promoting translation or overexpression of PD-L1 in ET macrophages.

To test the hypothesis, we performed a similar T cell proliferation assay^44^ to determine the effect of either G9a ko or METTL3 ko on T cell function under the TL/ET condition. We compared T-cell activation and proliferation by incubation of wild type, G9a ko, or METTL3 ko Raw cells collected in N, NL, and TL with T cells from a P14 transgenic mouse. By monitoring multiple markers of T-cell activation, including CD25, CD44, and CD69, we observed that the co-existing TL/ET wild type cells suppressed activation of CD8 T cells, whereas incubation with G9a ko or METTL3 ko cells produced efficiently activated T cells (**Fig. 2e**, upper panel). Similarly, we found a greater number of proliferated P14 CD8 T cells after six days of co-incubation of T cells with either G9a ko or METTL3 ko, whereas wild type cells lost the ability to promote proliferation of T cells in ET (**Fig. 2e**, lower panel). In line with downregulation of proteins that were highly enriched in T cell activation in severe COVID-19 patients, these results showed that G9a and METTL3 impair T cell function by promoting overexpression of immune checkpoint proteins such as PD-L1.

### Lysine methylation by G9a is critical for the stability of METTL3 complexes in ET macrophage gene-specific translation

We next investigated the function of the G9a-METTL3 interaction in ET macrophages. Briefly, we co-transfected equal amounts of HA-tagged G9a and Flag-tagged METTL3 into the LPS-responsive, TLR4/CD14/MD2 293 cells^45^ under N, NL, or TL. Notably, both METTL3 and G9a (input) showed significantly increased co-expression specifically in T or TL (ET). Immunoprecipitation with anti-HA antibody confirmed TL-specific interaction of the two proteins. Importantly, the METTL3-G9a interactions also increased with prolonged LPS stimulation that mimicked ET (**Fig. 3a**). Notably, the composition of the UNC0965-captured G9a interactome from ET/TL macrophages showed significant overlap with the composition of the G9a lysine methylation (Kme) proteome^46,47^; eight nonhistone G9a substrates were identified as ET-specific G9a interactors. These results indicated that certain G9a interactors are ET-specific G9a substrates.

**Fig.3.**
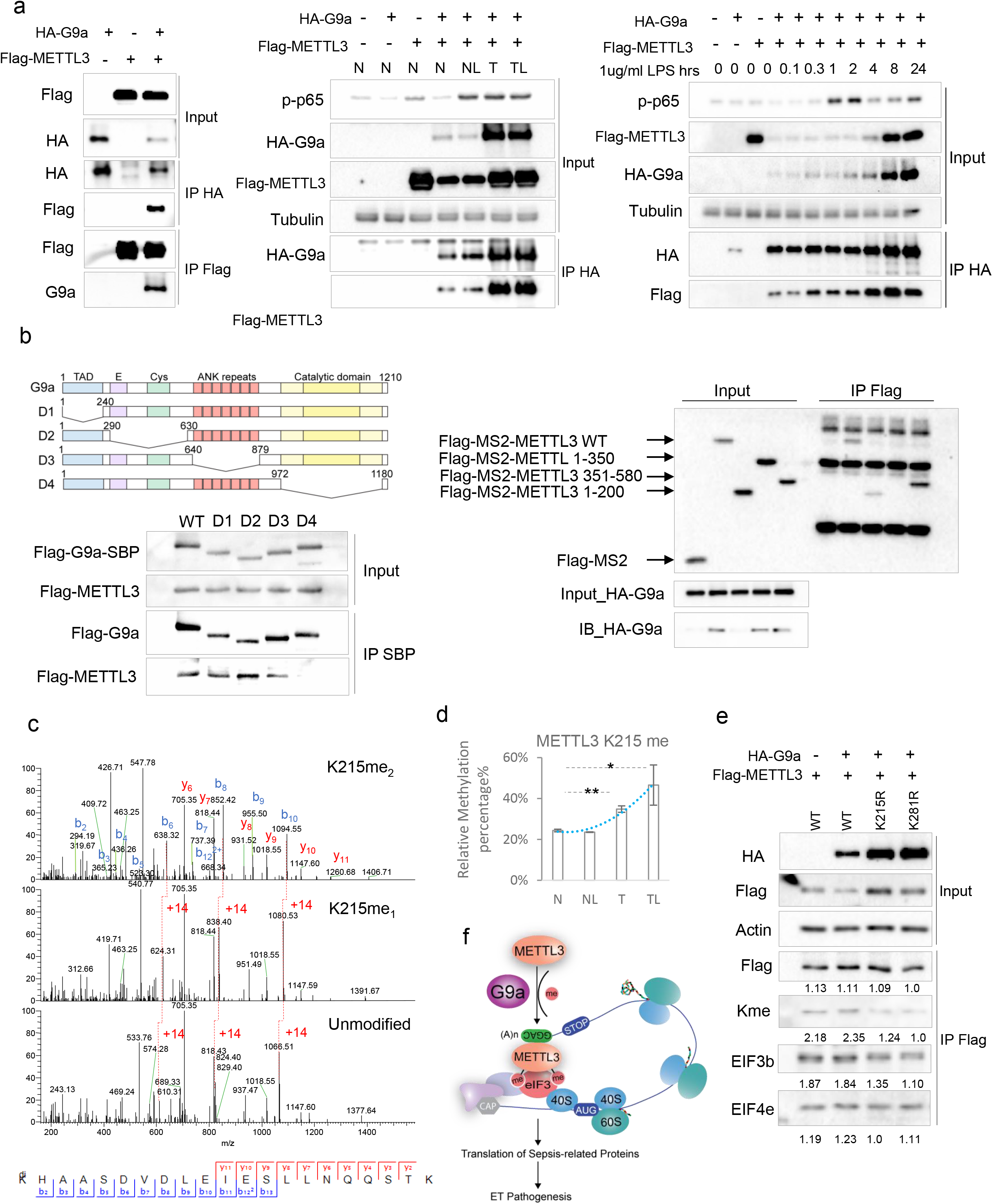
G9a-mediated lysine methylation is implicated in the METTL3-mediated translation in ET macrophages. **a**. (Left): G9a interacts with METTL3, the interactions were confirmed by both ‘forward’ HA-G9a IP and ‘reverse’ Flag-METTL3 IP; (middle): METTL3 specifically interacts with chronically active G9a in LPS-tolerant 293-TLR4/CD14/MD2 cells. Cell responses were monitored by p65 phosphorylation; (right): Timecourse dependent interactions between HA-G9a and Flag-METTL3 under N, NL, T, or TL. **b**. Mapping the interacting domains of G9a and METTL3; (Left): Lacking the catalytic domain of G9a (CD) disrupts the interaction with METTL3; (right): C-terminus deletion (201-580) mutant of METTL3 (Flag-MS2-METTL3 1-200) abolished the interaction with G9a. **c**. G9a methylates multiple METTL3 lysine in ET. MS/MS spectra of the tryptic peptides of METTL3 bearing mono-Kme or di-Kme. b- or y-ions were labeled in blue and red, a methylated ion has a 14-dalton mass shift compared with its unmodified counterpart. **d**. LPS-induced methylation dynamics of METTL3 under different inflammatory conditions. Flag-METTL3 and HA-G9a was transiently transfected into 293-TLR4/CD14/MD2 cells which then were subjected to LPS stimulation (N, NL, T, TL) 24 h after transfection. LFQ was based on relative peak areas of the identified methylated peptides and corresponding nonmethylated counterpart. Error bars show the standard deviation from two independent experiments each with duplicates. Asterisks indicate the statistical significance: **P□<□0.01, *P□<□0.05. **e**. Removal of lysine methylation weakened METTL3 interactions with eIF3b. ‘WT −wild type METTL3; ‘K215R, K281R’ – K-to-R mutant of METTL3. Interaction strength was determined by densitometry.

Considering that some G9a interactors are ET-specific G9a substrates, we examined whether the ET-specific interaction between G9a and METTL3 implied that G9a methylates METTL3 in ET. With domain-truncated versions, we found that the catalytic domain of G9a interacts with the C-terminus (aa 201-580) of METTL3 (**Fig. 3b**). We then immunoprecipitated Flag-METTL3 from the cotransfected TLR4/CD14/MD2 293 cells in TL; immunoblotting with an anti-mono- or di-methylysine antibody showed that METTL3 was methylated (**Extended Data Fig. 3**). We then used LC-MS/MS to sequence the tryptic digests of Flag-METTL3 and unambiguously identified four mono- or di-methyl-lysine (Kme) sites in the C-terminus of METTL3 that directly interact with the G9a catalytic domain (**Fig. 3c**). In addition, using LFQ MS to measure the differences in these Kme sites in different inflammatory conditions, we identified one Kme site (K215) that showed increased abundance in ET (**Fig. 3d**), thereby confirming that these METTL3 lysines are targets of constitutively active G9a. Du et al. reported that SUMOylation of METTL3 at lysine 215 modulates the RNA N^6^-adenosine-methyltransferase activity of METTL3.^48^ Also, E3 SUMO-protein ligases PIAS1, RanBP2, and CBX4, and a SUMO1-specific protease SENP1, were identified as ET-specific interactors of METTL3. SENP1 reduced METTL3 SUMOylation at multiple lysine sites including K215.^48^ These results confirmed the propensity of METTL3 to be post-translationally modified and the functional relevance of distinct modification (e.g., lysine methylation) types in ET macrophages. Notably, the transient nature of enzyme-substrate interaction probably explained why ChaC-MS did not unambiguously identify the endogenous G9a-METTL3 interaction.

Initiation is usually the rate-limiting step in translation, and METTL3 enhances translation of target mRNAs by recruiting eIF3 to the initiation complex.^20^ We made^49^ single and double, nonmethylatable mutants of METTL3, i.e., K215, K281, and K327-to-R (KXR), and used immunoblotting to compare the strength of binding of indicated proteins to the Flag-METTL3 versus Flag-K215R or K281R METTL3 mutants in the ET TLR4/CD14/MD2 293 cells. We observed that Kme absence (confirmed with Kme antibodies) weakened METTL3 interaction with eIF3 by at least 30% (**Fig. 3e**), a deficiency that was expected to impair METTL3-mediated translation.^20^ Notably, the METTL3 interaction with the 5’ cap-bound eIF4e was unaffected by the K215R or K281R mutation, a finding that agreed with the fact that METTL3 is critical for 5’UTR m^6^A mRNA to promote cap-independent translation.^50^ A mechanism underlying G9a methylation-activated, METTL3-mediated translation (**Fig. 3f**) is postulated based on results from the abovementioned combined approaches.

### G9a coordinates a widespread acceleration of gene-specific translation in ET

ChaC-MS identifications of additional translation regulators as G9a interactors indicated that the G9a-METTL3 axis is only one part of a translation regulatory network involving G9a (**Fig. 1a**). Thus, we employed our improved of AACT^51^ or SILAC pulse-labeling translatome^52^ strategy (**Fig. 4a** and **Supplementary Methods**) to profile, in a nonbiased proteome-wide manner, all genes that showed G9a-dependent translation, i.e., the ‘G9a-translated proteins’. Accordingly, we determined individual protein rates of synthesis, degradation, and overall turnover in snapshot samples collected at different times from cultures of wild type, G9a ko, and UNC0642^53,54^-treated macrophages. Here, genetic (G9a ko) and pharmacologic inhibition (UNC0642 treatment) of G9a decreased synthesis or turnover of G9a-translated proteins.

**Fig.4.**
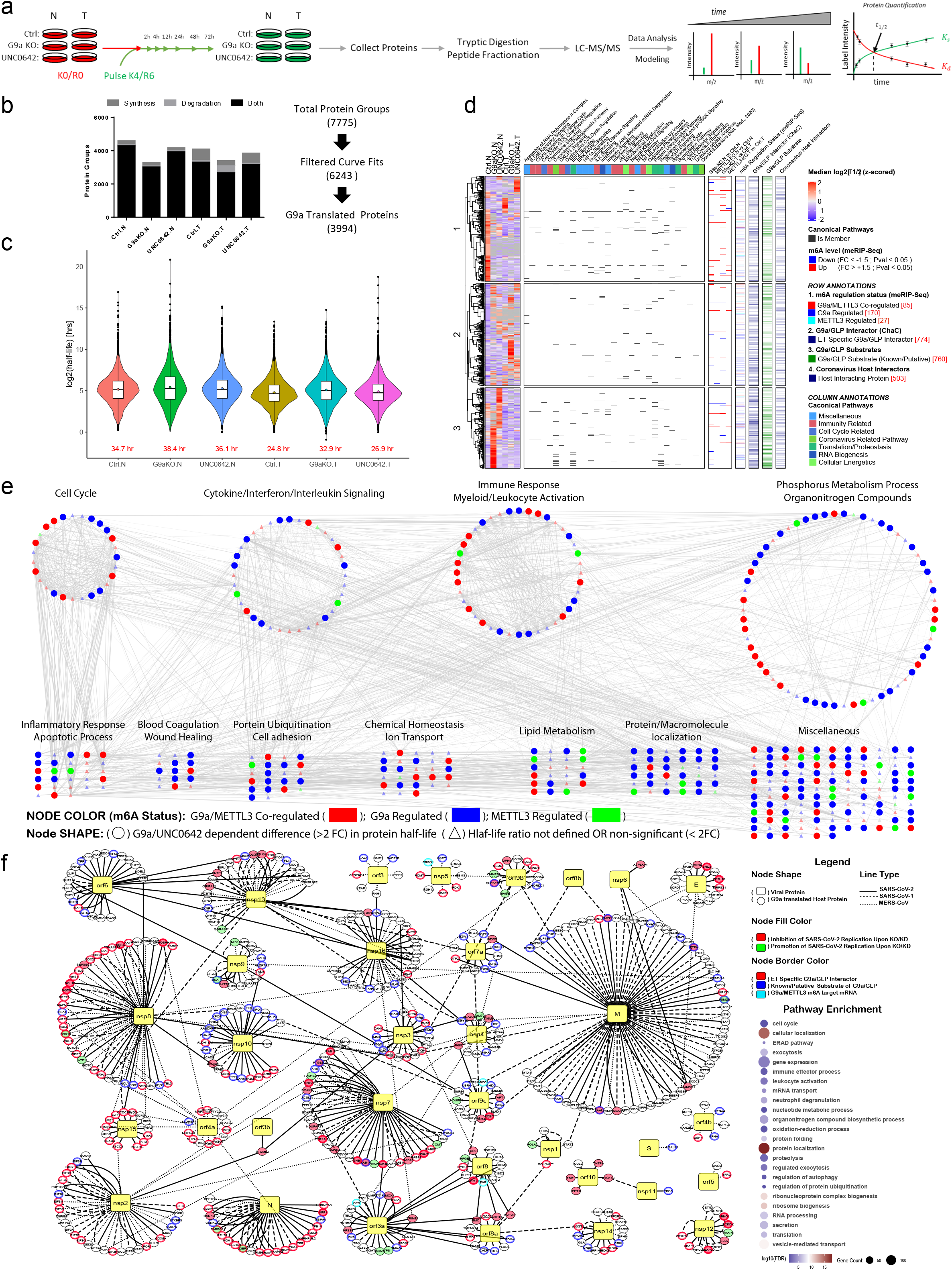
Constitutively active G9a promotes translation of SARS-CoV-2-evolving pathway components. **a**. Schematic of AACT-pulse labeling translatome strategy for determining the rates of protein synthesis, degradation, and turnover in N and TL/ET conditions. Raw macrophages grown in K0/R0 containing medium were pulse-labeled with K4/R6, with or without UNC0642 treatment, and harvested at 2h. 4h, 8h, 24h, 48h, and 72h. Decreasing and increasing K4/R6 labels were used to determine rates of protein synthesis, degradation, and turnover (see **Supplementary Methods** for a detailed explanation of the curve fitting). **b**. Numbers of the proteins showing >2 fold-change (FC) differences in half-life upon G9a-KO or UNC0642 treatment compared with wild type cells. **c**. The wild type macrophages in ET/T had faster turnover (shorter median half-life). Distribution of protein turnover (*log2*[*t*_12_]) is depicted by violin plots; sample median (horizontal line), mean (diamond shape) and IQR are shown by overlaid boxplot. Red text shows median protein turnover/half-life [in hours]. **d**. Hierarchical clustering of G9a-translated genes (mRNAs) and associated pathways. Certain G9a-translated proteins were identified as G9a/GLP interactors, non-histone substrates, or with m^6^A. Translation of numerous COVID-19 markers, SARS-CoV2 host interactors and other coronavirus-related proteins were also affected by G9a ko or inhibition in T. **e**. Protein interaction networks for G9a/METTL3 regulated m^6^A-target genes shows that G9a ko or inhibition alters protein translation dynamics for ~59.7% of the proteins. Representative pathways from each cluster are also shown. **f**. Virus-host protein-protein interaction map depicting G9a-translated proteins identified by our translatome strategy. Human proteins are shown as circles, whereas viral proteins are represented by yellow squares. Each edge represents an interaction between a human and a SARS-CoV-2 (solid line), SARS-CoV-1 (dashed line) or MERS-CoV (dotted line) protein with several interactions shared between these three viruses. Node border colors show that several of the G9a-translated host proteins are ET-specific G9a interactors (red), G9a/GLP substrates (blue) or G9a/METTL3-regulated m^6^A mRNAs (cyan). Node fill color shows that genetic perturbation of several of these G9a-translated proteins (red) SARS-CoV-2 replication/infection with some showing the opposite effect (green). Pathway enrichment scores for 503 G9a-translated proteins are shown on the side. ET-specific G9a interactors [ChaC] and G9a/METTL3-regulated m^6^A targets [MeRIP-Seq] were defined in this study, whereas host-virus physical interactions, effect of genetic perturbation on SARS-CoV-2 replication/infection, and G9a/GLP substrate definitions were curated manually from literature sources. (also see **Extended Data Table 3b**).

Briefly, cells grown with Lys0-Arg0 (K0/R0, ‘light’, L) were pulse-labeled with Lys4-Arg6 (K4/R6, ‘medium’, M) supplemented media. At 2h, 4h, 8h, 24h, 48h, and 72h after the media switch, proteins extracted from harvested cells were subjected to tryptic digestion, fractionation, and LC-MS/MS. This experimental design yields (i) increasing signals from the K4R6-labeled protein molecules due to nascent protein synthesis, and (ii) decreasing signals from K0R0-labeled proteins due to degradation or secretion of pre-existing protein molecules. Accordingly, the inhibitor-induced rate changes in nascent protein synthesis or protein degradation were quantified by the intensities of L or M labels at different time points in wild type, G9a ko, and UNC0642-treated macrophages, respectively, under non-stimulated (N) or prolonged endotoxin stimulation (T) conditions (**Fig. 4a)**. The model assumed steady-state equilibrium conditions, in which the rate of increase was counterbalanced by the rate of decrease, leading to stable intracellular protein levels. Effects of amino acid recycling and differences in cell division rate between different conditions were also considered. Fit qualities were estimated using least-squared regression (*R*^2^), root-mean-squared error (RMSE), with additional thresholds on fitted parameters to ensure good/meaningful estimates of protein turnover (**Extended Data Table 3a**).

We obtained synthesis and/or degradation half-lives for 6,243 protein groups in the combined dataset (i.e., wild-type/Ctrl, G9a-KO, UNC0642-treated cells under N and T conditions). For 78–94% of fitted proteins, information was available for both AACT-label increase and decrease, which provided an internal duplicate measurement of protein turnover time for each sample in a steady-state system (**Fig. 4b** and **Extended Data Table 3b**). Half-life determination was reliable across labeling pairs (R = 0.99) and cell culture replicates (R = 0.29-0.99) as evidenced by high Pearson’s correlation coefficients and covariance (**Extended Data Fig. 4a & 4b**). Under N and T conditions, whereas the principle component analysis hinted towards distinct proteostasis landscapes in macrophages, G9a knock-out or inhibition produced large effects on protein turnover compared with wild type macrophages (**Extended Data Fig. 4c**). Consequently, we observed significant pairwise differences in global protein turnover time upon G9a inhibition as well as ET (**Extended Data Fig. 4d**). Estimated protein turnover times spanned four orders of magnitude (with some outliers), from minutes to thousands of hours. Similar to SARS-CoV-2-upregulated global translation in multiple organs of severe patients, ET also increased the rates of global translation or protein turnover as evidenced by shorter median half-lives in T (24.8-32.9 h) compared with N (34.7-38.4 h). The ET-accelerated translation rates were reduced in G9a ko or UNC0642-treated macrophages specifically under T (32.9, 26.9h) (**Fig. 4c**). In agreement with the polysome results that showed G9a/METTL3-dependent protein synthesis (**Fig. 1c**), this nonbiased translatome profiling indicated that constitutively active G9a accelerated global protein synthesis and degradation (i.e., global proteostasis) in ET.

### Constitutively active G9a promotes translation of specific protein components in SARS-CoV-2 pathologic pathways related to the host response and viral replication

Among 6,243 proteins with AACT-pulse labeling-measured turnover rates, 3,994 were identified as G9a-translated proteins based on pairwise comparisons of protein half-lives (**Extended Data Fig. 4e**). Pathway enrichment analysis indicated that all G9a-translated proteins are primarily involved in immune responses involving B-cell, T-cell, NK-cell, chemokine, interferon, interleukin signaling, G1/S checkpoint and cyclin signaling, RNA biogenesis such as splicing and mRNA degradation, RNA Pol II assembly, translation/proteostasis including EIF2/4, ubiquitination, SUMOylation, unfolded protein response signaling, cellular energetics including oxidative phosphorylation and TCA cycle signaling, and coronavirus-related pathways. These results showed that, coincident with ET-phenotypic increases in global rates of protein turnover (see **Fig. 4c**), constitutively active G9a upregulates diverse pathways by activating the translation of particular pathway genes (**Extended Data Fig. 4f**). A closer look at net protein turnover in individual pathways indicated that the global protein turnover trend did not completely capture the underlying complexity of G9a/ET regulated proteostasis because the effects of G9a inhibition on individual pathways varied greatly (**Extended Data Figs. 5**). **Fig. 4d** shows a summary heatmap depicting median protein half-lives of G9a-translated proteins and their associated pathways in ET macrophages. Interestingly, of these 3,994 G9a-translated proteins, 774 proteins were identified by UNC0965 ChaC-MS as ET-phenotypic G9a interactors, and 760 proteins are known or putative substrates^55,56^ of G9a/GLP **(Extended Data Table 3b**). G9a mediated methylation of a nonhistone substrate FOXO1 has already been shown to induce proteasomal degradation^57^. Therefore, our results not only support our identification of METTL3 as both G9a interactor and a nonhistone substrate of G9a (**Fig. 3c**) but also indicate that G9a may upregulate gene-specific turnover by interacting with or methylating select translation regulators. Notably, 472 G9a-translated proteins were encoded by G9a and/or METTL3-regulated m^6^A mRNAs (**Fig. 4d** and **Extended Data Table 3b**), of which 282 proteins (~59.7%) showed more than two-fold difference in turnover time in a G9a-dependent manner (**Fig. 4e**). These m^6^A mRNA-encoded, G9a-translated proteins are associated with cell cycle, cytokine/chemokine/interleukin signaling, myeloid/leukocyte activation, blood coagulation and wound healing, proteostasis including ubiquitination, localization (i.e. endocytosis, vesicle transport) signaling, organo-nitrogen metabolism, i.e., lipid, nucleotide, amino-acid synthesis, and ion transport **(Fig. 4e)**. Thus, combined results from ChaC-MS, m^6^ARIP-Seq, and translatome analysis validated the regulatory function of the G9a-METTL3-m^6^A axis in ET-phenotypic translation activation, which accounted for a subset of G9a-translated genes (472 out of 3994). Clearly, *via* interactions with distinct translation regulators other than METTL3 (**Fig. 1b**), G9a coordinates additional, as yet unknown, mechanisms to facilitate genespecific translation (**Fig. 4d**).

Strikingly, almost all of the G9a-translated pathways that we identified by translatome profiling (**Figs 4d and 4e**) have been implicated in SARS-CoV-2 life cycle and COVID-19 pathogenesis ^4,8,10,15,19,58,59^. Indeed, we observed G9a-dependent turnover for 11 COVID-19 markers ^8,58^ (**Extended Data Figs. 4g-4h**), 503 SARS-CoV-1/2 & MERS-CoV host interactors ^29,30^ (**Fig. 4f**) and 66 other coronavirus pathogenesis pathway-related proteins (**Extended Data Fig 5b**). Several of these G9a-translated proteins were identified as ET-specific G9a interactors, nonhistone G9a substrates or G9a/METTL3-dependent m^6^A targets (**Extended Data Fig. 5b**; also see **Extended Data Table 3b**), which supported the translation regulatory function of G9a in COVID-19 pathogenesis. More importantly, genetic perturbation of several of these G9a translated host interactors adversely affected SARS-CoV-2 replication and infection ^13,30^ (**Fig. 4f)**. Similarly, several host factors critical for SARS-CoV-2 infection identified in siRNA/CRISPR based screens are closely related to G9a complex ^13,30^ (**Extended Data Fig. 7**). In line with COVID-19 hallmarks of a systemic cytokine storm, excessive infiltration of monocytes, dysregulated macrophages, and impaired T cells^4^, we observed faster turnover of proteins that belong to immune response pathways involving B-cell receptor, T-cell receptor, NK-cell, chemokine, interferon, interleukin, Jak/Stat, NF-kB and CXCR4 signaling. Further, pharmacologic inhibition of G9a downregulated these pathways by reducing the translation/turnover rates of major pathway components (**Extended Data Fig. 5c**). Similarly, proteins involved in splicing, unfolded protein response, and translation initiation/elongation were upregulated following SARS-CoV-2 infection ^19^. Consistent with these findings, we observed increased turnover for >150 proteins related to translation/proteostasis including EIF2/4, unfolded protein response, SUMOylation and ubiquitination signaling and RNA biogenesis such as spliceosomal cycle and RNA degradation pathways in ET macrophages, whereas G9a inhibition reversed these effects by reducing their turnover times (**Extended Data Fig. 5e** & **5g**). In short, constitutively active G9a regulates specific genes at the translational or posttranslational level to drive ET-related, SARS-Cov-2-induced pathogenesis, and inhibition of G9a and its associated proteins hinders coronavirus replication and infection. Thus, G9a and its associated proteins are potential drug targets to treat COVID-19 and other coronavirus-related ailments, and these targets merit further molecular and clinical study.

### Enzymatic inhibition of G9a or Ezh2 similarly mitigated overexpression of COVID-19-characteristic proteins

Interestingly, we also identified as ET-specific interactors of G9a the histone methyltransferase Ezh2 (Enhancer of zeste homolog 2) and two Polycomb Repressive Complex 2 (PRC2) components, SUZ12 and EED (**Fig. 1b**). Further, we were intrigued by the G9a-dependent translation of the PRC2 complex components, which includes Ezh2, in ET (**Extended Data Fig. 6c**). Like the elevated level of G9a mRNA, most PRC2 complex components were overexpressed in COVID-19 patients with high viral load (**Extended Data Figs. 6a & 6b**). Because there are clinically validated inhibitors of Ezh2 in antitumor clinical trials, we considered Ezh2 inhibitors for severe COVID-19 therapy and compared the proteomic effects of an Ezh2 inhibitor (UNC1999)^60^ with a G9a inhibitor. We used various quantitative proteomic approaches^24–26^, i.e., a multiplex TMT quantitative proteomic method to analyze the intracellular proteins collected from the pellets of N, NL, and T Raw macrophages, with or without inhibitor treatment. In parallel, LFQ proteomic approach was used to comparatively identify the proteins whose secretion showed dependence on the treatment by either G9a or Ezh2 inhibitor or both, respectively. Based on their overexpression in ET macrophages, we identified 43 proteins (**Fig. 5a**) whose abundances were suppressed by either or both inhibitors. Strikingly, >80% of these proteins that showed G9a- or Ezh2-dependent overexpression in ET macrophages were associated with ARDS-related cytokine storm or clinical ARDS, with some proteins found in the sera of severe COVID patients^6^ (**Fig. 5b and Extended Data Table 4**).

**Fig.5.**
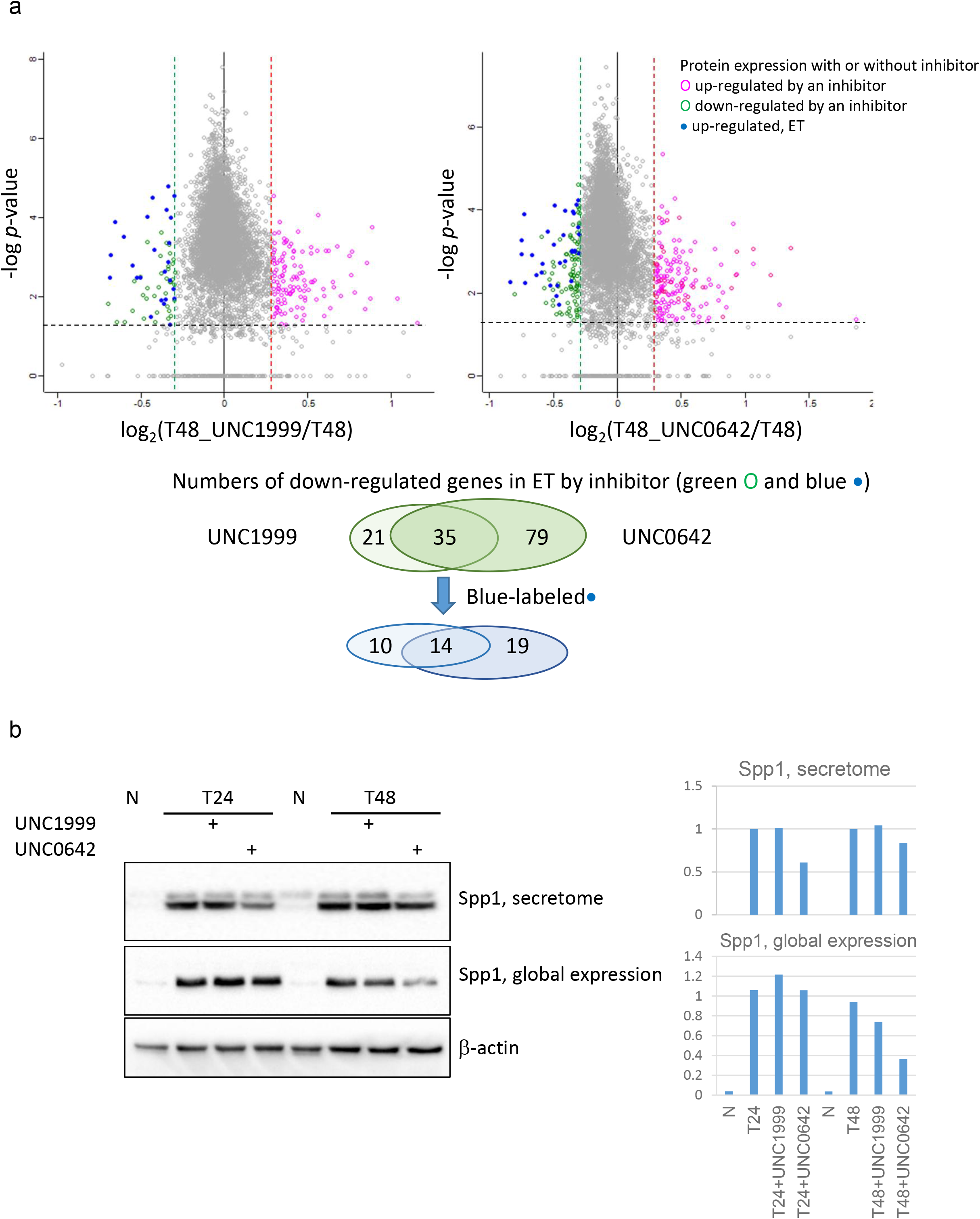
The enzymatic inhibition of G9a or Ezh2 mitigated overexpression or secretion of COVID-19-characteristic proteins. **a**. Differentially expressed proteins induced by G9a- or Ezh2-inhibitor at ET condition. T-test p-values were generated from three technical replicates. Proteins with log2 (fold-change) beyond 0.30 or below −0.30 with p value lower than 0.05 were considered as significantly differential expression. Number of significantly down- (green and blue) and up- (red) regulated proteins were shown on lower panel. Proteins labeled in blue were up-regulated in ET cells compared to naïve cells (N). **b**. Immunoblot analysis of G9a- or Ezh2-inhibitory changes of SPP1, a COVID-19-characteristic protein.

## DISCUSSION

### Discovery of nonepigenetic function of G9a in driving SARS-CoV-2 immunopathogenesis

In 10-20% of patients, SARS-CoV-2 infection progresses to ARDS or/and severe pneumonia or multi-organ damage/failure; most of these patients are vulnerable because of pre-existing chronic conditions. The hyperinflammatory response mediated by SARS-CoV-2-dysregulated macrophage activation results in a cytokine storm, which is the major cause of disease severity and death.^6^ Also, the immunologic characteristics of COVID-19, especially in severe cases that require ICU care, include reduced counts of CD3+, CD4+ and CD8+ T lymphocytes or lymphopenia,^61^ and significantly increased levels of certain serum cytokines or hyperinflammation. In addition, as indicated by significantly higher levels of a T cell exhaustion marker PD-1, the surviving T cells in severe patients appeared to be functionally exhausted. We discovered a new gene-specific translation mechanism of hyperinflammation, lymphopenia and viral replication in which the constitutively active G9a/GLP interactome coordinates the networked, SARS-CoV-2-dysregulated pathways that determine COVID-19 severity.

Endotoxin-tolerant macrophages have molecular characteristics similar to chronic inflammation-associated complications (e.g., ARDS and sepsis), including downregulation of inflammatory mediators and upregulation of other antimicrobial factors. These complications systemically contribute to impaired adaptive immunity (e.g., T-cell function impairment or a poor switch to the adaptive response) and susceptibility to secondary infection with an organ-damaging cytokine storm.^11^ The current model for inflammation control in ET macrophages was derived from mRNA expression studies.^62^ Foster et al. reported that the chromatin modification landscape was differentially programmed in a gene-specific (pro-inflammatory versus antimicrobial genes) manner. That is, with prolonged LPS stimulation, the promoter chromatin of pro-inflammatory genes transitioned from transcriptionally activate to transcriptionally silent and LPS-induced transcripts regulated increased expression of other antimicrobial genes in ET.^62^ In parallel with G9a’s canonical epigenetic function for transcriptional silencing of pro-inflammatory genes, particular G9a interactors associated with chromatin regulation such as BRD2/4 were found to affect SARS-CoV-2 replication^30^. Additional G9a interactors including the components of chromatin remodeling complexes such as SMARCA2/4, SMARCC2, CREBBP, were characterized as proviral host factors essential for SARS-CoV-2 infection^13^ (**Fig. 1c**). Conversely, upon restimulation, antimicrobial proteins are unlikely to be derived from time-consuming transcription processes as the turnover of mRNAs is an energy-cost-effective process to respond to an infection, which suggests that overexpression of antimicrobial genes in ET is regulated by under-characterized translation mechanisms. Coincident with its constitutive activity in ET^14^ and overexpression in COVID-19 patients with high viral load^15^, here G9a is characterized as a noncanonical (nonepigenetic) regulator of genespecific translation to drive SARS-CoV-2 immunopathogenesis. Mechanistically, via ET-phenotypic interactions with METTL3, a known gene-specific translation activator, and other translation regulators, G9a activates diverse translation regulatory pathways associated with major clinical characteristics of COVID-19. Specifically, on the basis of results from MS/MS-sequencing and biochemical assays, we clarified the promoting activity of the G9a-METTL3 axis in gene-specific translation, in which G9a-mediated methylation enhances the stability of the METTL3/m^6^A-mediated translation regulatory complex (**Fig. 3f**).

From a broad view, SARS-CoV-2 infection upregulates G9a/GLP (**Extended Data Fig. 1a**) which ‘reshapes’ the translation regulatory pathways and, in turn, activates the translation of a range of genes for SARS-CoV-2 immunopathogenesis. Briefly, Bojkova et al.^19^ reported that SARS-CoV-2 infection led to increased expression of proteins associated with translation initiation and elongation, alternative splicing, mRNA processing, and nucleic acid metabolism. Likewise, in the ET macrophages that showed similar immunologic features to severe COVID-19, we found that most of these SARS-CoV-2-upregulated translation components in infected cells were ET-phenotypic interactors and nonhistone substrates of constitutively active G9a (**Fig. 4**). Further, by nonbiased screening of the ET macrophage translatome with time-dependent inhibition of G9a, we identified genes that are translated at higher rates in a G9a-dependent manner as ‘G9a-translated proteins’. Clustered by functional pathway enrichment, ~4,000 G9a-translated proteins were merged into two major networks associated, respectively, with the host response to SARS-CoV-2 infection (**Fig. 4e**) or host-virus interactions (**Fig. 4f**). For example, COVID-19 severity was linked to multiple SARS-CoV-2-induced, G9a-translated host response pathways, primarily associated with immune complement and coagulation dysregulation^15^, IFNs- and IL-6-dependent inflammatory responses^63^, and ER-associated degradation (ERAD)^64^ (**Fig. 4e**). The G9a-translated components of complement and coagulation pathways including C5aR1, SERPINE1, CR1L were upregulated in severe patients^65^. Overexpression of the C5a-C5aR1 axis was associated with ARDS in COVID-19 patients^66^. Specifically, increased PD-L1 levels in monocytes and dendritic cells ^67^ and elevated levels of C5aR1 in blood and pulmonary myeloid cells ^59^ contribute to COVID-19-characteristic hyperinflammation and ARDS. G9a inhibition reduced the turnover rates of both proteins in ET macrophages (**Extended Data Fig. 4h**). These COVID-19-associated networks composed of G9a-translated proteins provide a compelling rationale to suspect that G9a inhibition will adversely affect SARS-CoV-2-upregulated pathways associated with not only the impaired host response but also with viral infection and replication.

### Inhibitor mechanism of action for severe COVID-19 therapeutics

Once immunopathologic complications such as ARDS or pneumonia occur, particularly in patients with pre-existing chronic inflammatory diseases, antiviral treatment alone is less effective and should be combined with appropriate antiinflammatory treatment. However, current immunomodulatory measures failed to improve prognosis of patients vulnerable to a cytokine storm.^68^ Our ChaC-MS dissection of G9a-associated pathways revealed a novel mechanism of inhibiting G9a for COVID-19 therapy. By ET-phenotypic interactions with primary components of the SARS-CoV-2 upregulated translation machinery, the enzymatic activity of G9a affects multiple inter-promotional pathways associated with COVID-19-characteristic hyperinflammation, lymphopenia, and virus replication. Accordingly, global translatome profiling of G9a inhibitor-treated ET macrophages identified a profile of G9a-dependent protein overexpression similar to the systemic cytokine profiles observed in COVID-19 patients. Specifically, inhibition of G9a enzymatic activity reduced the expression of eleven ARDS- or sepsis-related proteins: SPP1, CCL2, IL1RN, CXCL2, SQSTM1, ANPEP, PLAU, PELI1, PROCR, DST, and FABP4. These proteins had higher abundances in severe versus mild patients, or mild patients compared with healthy individuals.^6^ Thus, unlike most anti-inflammatory therapies that target only one cytokine/chemokine at a time, G9a-targeted therapy can suppress the systemic hyperinflammatory response by simultaneous inhibition of multiple components of a COVID-19 cytokine storm. In parallel, G9a depletion can also restore T cell function and reverse lymphopenia in COVID-19 patients. Because ChaC-MS identified Rps14 and SF3B1, the anti-SARS-CoV-2 targets^19^, as ET-phenotypic G9a interactors, G9a-targeted therapy can be combined with Rps14- or SF3B1-inhibition antiviral therapy to improve the efficacy of single target therapy. Also, because an Ezh2 inhibitor showed inhibitory effects similar to the effects of a G9a inhibitor, the Ezh2 inhibitors with proven safety for cancer therapy can be repurposed for COVID-19 therapy.

## CONCLUSION

The ability to develop targeted therapies to minimize mortality of severe patients depends on a detailed understanding of SARS-CoV-2-dysregulated chronic inflammatory pathways that determine COVID-19 severity. Our results from chemoproteomics, MeRIP-seq, translatome proteomics, and molecular/cell biology revealed a novel mechanism of inhibitor action on COVID-19. Specifically, via ET-phenotypic G9a-interacting translation machinery, SARS-CoV-2 may evolve a G9a-associated mechanism of gene-specific translation activation to modulate host response, evade the host immune system, and promote viral replication and infection. In endotoxin-tolerant macrophages that share similar immunopathologic characteristics with SARS-CoV-2 dysregulated macrophages, we dissected the G9a-associated pathways that cause individuals with preexisting chronic inflammatory diseases to be highly susceptible to secondary infection by SARS-CoV-2. Specifically, we discovered a widespread translational function of constitutively active G9a in the impairment/depletion of T cell function and the production of organ-damaging cytokine storm. Importantly, combined results from COVID-19 patients with pre-existing conditions, genome-wide CRISPR screening, and CoV-host interactome mapping validated our mechanistic findings in ET and identified numerous G9a interactors or G9a-translated proteins in the interconnected networks associated, respectively, with the host response to SARS-CoV-2 infection or host-virus interactions. Compared with current trials focused on either antiviral or anti-inflammatory therapy, this single-target inhibition of G9a-associated pathways was responsible for multifaceted, systematic effects, namely, restoration of T cell function to overcome lymphopenia, mitigation of hyperinflammation, and suppression of viral replication. Our studies have paved new roads that combine immunomodulatory and antiviral therapies with more effective COVID-19 therapies with minimal side effects. Importantly, because both G9a and its complex components were overexpressed in infected host cells by other CoV strains (e.g., SARS-CoV-1 and MERS-CoV^30^), our G9a-targeted therapy is refractory to complications induced by emerging antiviral-resistant mutants of SARS-CoV-2, or any virus, that hijacks host responses.

## ACKNOWLEDGEMENTS

This work was supported in part by grants NIH R01 GM133107-01 and UNC University Cancer Research Fund (UCRF) (to X.C.), and R01GM122749, R01HD088626 and an endowed professorship from the Icahn School of Medicine at Mount Sinai (to J.J.), and R01GM102413 (to J.G.C). This work used Thermo-Fisher Scientific Q Exactive HF-x that was upgraded with funding from UNC Lineberger Comprehensive Cancer Center (LCCC). This work also used the AVANCE NEO 600 MHz NMR Spectrometer System that was upgraded with funding from a National Institutes of Health SIG grant 1S10OD025132-01A1. We thank Dr. Howard Fried for editorial assistance. This invention of the mechanism of inhibitor action is protected by United States Provisional Patent Application that was filed by the University of North Carolina-Chapel Hill (UNC 21-0060). The new indication of using clinically trialed Ezh2 inhibitors for COVID19 therapy is protected by US provisional patent application #63/113,211 as ‘Use of Method’. The clinical trial application is under consideration with the FDA Coronavirus Treatment Acceleration Program (CTAP).

## AUTHOR CONTRIBUTIONS

L. W. performed ChaC-MS and translatome experiments, biological and cell biology assays, and wrote the report. A.M. developed the software for translatome analysis, performed network analysis of clinical data, and wrote the report. L.X. and E. H. F. performed inhibitor treatment, sample preparation and processing for MS/MS experimental analysis, and analyzed data. F. Z., J. S., and H.S. performed RNA-seq, MeRIP-seq, and analyzed the data. B.W. and Y.Y.W analyzed T cell activation/proliferation. L. M. and J.G.C performed cell cycle data collection and analysis. E.M.L. and N.J.M performed polysome experiments, and analyzed data. C. W., C. G., and K. T. assisted with clinical analysis. Y.X. and J.J. provided UNC0965, UNC0642, and UNC1999. X.C. conceived and designed the project and experiments, analyzed and interpreted data, and wrote the manuscript.

## METHODS

### Chemicals and reagents

Cell culture media, other components, and fetal bovine serum were obtained from Gibco. Trypsin was purchased from Promega. LPS (Escherichia coli 0111:B4, ultrapure) was purchased from InvivoGen (USA, Invivogen, cat# tlrl-pelps). HEK 293 stable TLR4-MD2-CD14 cell line was also purchased from InvivoGen. All chemicals were HPLC-grade unless specifically indicated. Raw264.7 and THP1 cells were purchased from ATCC (Manassas,VA). UNC0642 (G9a inhibitor) and UNC1999 (EZH2 inhibitor)^60^ were synthesized in Dr. Jian Jin’s lab. TMT 11-plex isobaric labeling reagent kit was purchased from Thermo Fisher (Cat. A34808). Antibodies against G9a (07-551) and H3K9me2 (07-441) were from Millipore; Antibodies against EIF4E(11149-1-AP), EIF3B(10319-1-AP), HNRNPA2B1(14813-1-AP),METTL3(15073-1-AP), YTHDF2(24744-1-AP),PD-L1 (66248-1-lg), CDC20(10252-1-AP), CDCA7(15249-1-AP), BIRC5(10508-1-AP), TPX2(11741-1-AP), TOP2A(24641-1-AP), NUSAP1(12024-1-AP), SPP1(22952-1-AP), SQSTM1(18420-1-AP), ACTIN(60008-1-lg) were from Proteintech. Antibody against p-p65 (S536)(3033) was from Cell Signaling. Antibody against Brg1 (H-88) (sc-10768x) was from Santa Cruz. Anti-HA (clone HA-7) and anti-flag M2 (clone M2) antibodies were from Sigma. m6A RNA methylation quantification kit was from Abcam(ab185912).

### Plasmids

Plasmid pHA-G9a was described^12^. pFlag-CMV2-METTL3, Plasmid pFlag-ms2-METTL3 and truncated expression plasmids were kindly provided by Dr. Richard I. Gregory^17^. Full-length pFlag-G9a-SBP and truncated plasmids (D1-D4) were kindly provided by Dr. Xiaochun Yu^69^.

### Cell Lines and Treatment

Raw264.7 cells were cultured in DMEM medium (Gibco). The human monocytic cell line THP-1 was maintained in RPMI 1640 medium (Gibco). All media were supplemented with 10% fetal bovine serum (Gibco), 100 U/ml penicillin and streptomycin. Cells were grown at 37 °C in humidified air with 5% carbon dioxide. For ChaC pull-down experiments, Raw264.7 cells were either unstimulated (‘N’) or subjected to a single LPS stimulation with 1 μg/ml (‘NL’) or first primed with 100 ng/ml LPS to induce endotoxin tolerance for 24h (‘T’), followed by a second LPS challenge at 1 mg/ml (‘TL’). For the condition requiring G9a inhibitor treatment, 1 μM UNC0642 was added at the time of cell plating.

For global protein profiling, Raw264.7 cells were pre-treated with 0.1 μg/ml LPS for 24 hours, then inhibitor UNC1999 (1 μM) or UNC0642 (1 μM) was added. The cells were collected after inhibitor treatment for 8 h, 24 h and 48 h.

For MeRIP-Seq, THP-1 cells were first incubated in the presence of 60 nM PMA overnight to differentiate into macrophages followed by 48 h resting in PMA-free medium. Cells were either left unstimulated (‘N’) or subjected to a single LPS stimulation at 1 μg/ml (‘NL’), or first primed with 100 ng/ml LPS to induce endotoxin tolerance for 24 h (‘T’), followed by the second LPS challenge at 1 μg/ml (‘TL’).

### UNC0965 pull-down and ChaC sample processing

Similar to our recent report,^16^ 1 mg nuclear protein extracted from Raw 264.7 macrophage cells was incubated overnight at 4°C with 2 nmole UNC0965 pre-coupled to 50 μl neutravidin-agarose (Thermo Fisher), and washed three times with 1 ml lysis buffer to remove non-specific bound proteins. For on-beads sampling and processing, five additional washes with 50 mM Tris-HCl pH 8.0, 150 mM NaCl were used to remove residual detergents. On-beads tryptic digestion was performed with 125 μl buffer containing 2 M urea, 50 mM Tris-HCl pH 8.0, 1 mM DTT, 500 ng trypsin (Promega) for 30 min at room temperature on a mixer (Eppendorf). The tryptic digests were eluted twice with a 100 μl elution buffer containing 2 M urea, 50 mM Tris-HCl pH 8.0, 5 mM iodoacetamide. Combined eluates were acidified with trifluoroacetic acid at final concentration of 1% (TFA, mass spec grade, Thermo Fisher) and desalted with a C18 stage tip.

### Sample preparation for TMT based global proteomic profiling

Cell pellets were resuspended in 8□M Urea, 50□mM Tris-HCl pH 8.0, reduced with dithiothreitol (5□mM final) for 30□min at room temperature, and alkylated with iodoacetamide (15□mM final) for 45□min in the dark at ambient temperature. Samples were diluted 4-fold with 25□mM Tris-HCl pH 8.0, 1□mM CaCl_2_ and digested with trypsin at 1:100 (w/w, trypsin : protein) ratio overnight at room temperature. Peptides were desalted on homemade C18 stage tips. Each peptide sample (100 μg) was labeled with 100 μg of TMT reagent following the optimal protocol^70^. The mixture of labeled peptides was desalted and fractionated into 12 fractions in 10 mM TMAB containing 5-40% acetonitrile.

### Polysome Fractionation

METTL3 KO, G9A KO, and Control Raw 264.7 cells were cultured in 10 cm plates until 80% confluent at the time of harvest. Cells were treated with 100 μg/ml cycloheximide (CHX; Sigma) for 10 min at 37^0^C. Media were removed and the cells were washed twice with 10 ml PBS containing 0.1 mg/ml CHX, scraped, pelleted by spinning for 10 min at 2200rpm at 4^0^C. After removing the supernatant, the cells were resuspended with 1ml lysis buffer (20 mM Tris-HCl, pH 7.4, 140 mM KCl, 5 mM MgCl_2_, 1% Triton X-100, 10 mM DTT) containing 0.1 mg/ml CHX and swelled on ice for 10 min followed by passing through a 27 gauge needle 5 times to break the cell membrane. Spin down the lysate at max speed for 10 min, the supernatant were carefully layered onto 10-50% sucrose gradients and centrifuged at 32,000rpm (no brake) in a Beckman SW-40 rotor for 2 h at 4^0^C. Gradients were fractionated and monitored by absorbance 254 nm.

### Transfection and CRISPR Knockout

293TLR4-MD2-CD14 cells were seeded in 6-well plates for 24 h before transfection, and the constructs were transfected or co-transfected into MCF7 cells using reagent jetPRIME (Polyplus). After 24 h, the cells were lysed directly in the plates by adding SDS-PAGE sample buffer, heating at 95°C for 5 min, and sonicating for 5 seconds to clear the lysate for immunoblotting.

For the CRISPR/Cas9 constructs, oligonucleotides for the sgRNA of human and mouse G9a and METTL3 as described below, were annealed and cloned in BsmBI-digested lentiCRISPRv2 (Addgene plasmid #52961). The empty vector was used as a negative control. Viral production was performed with a standard protocol. In brief, a total of 10 μg of plasmid, including target plasmid, pMD2.G (Addgene plasmid #12259) and psPAX2 (Addgene plasmid #12260) with a ratio of 10:5:9, was co-transfected into 293T cells with jetPRIME™. Viruscontaining media were collected 48 hours after transfection. MDA-MB-231 cells at 60-80% confluency were incubated with the virus containing media for 24-48 hours, and then subjected to 1.0 μg/mL puromycin selection. After 4-7 days puromycin selection, the stably transfected cells were collected for further analysis.

### RNA isolation and RT-PCR and m6A RNA methylation level

Total RNA was isolated using RNeasy kit (Qiagen). First-strand cDNA was synthesized by M-MLV reverse transcriptase (Promega, car# M170B) and diluted by a factor of 10 for quantitative PCR. Real-time PCR was performed using SYBR Green Master Mix (Thermo Fisher, cat# 0221). All measurements were normalized to GAPDH and represented as relative ratios. PCR primers for real-time PCR are summarized in Supplementary Table. The m6A level of isolated RNA was measured by using the m6A RNA methylation quantification kit.

### Cell Cycle and FACS Analysis

For flow cytometry analyses of cell cycle, cells were incubated with 10 uM EdU (Santa Cruz, sc-284628) for 30 minutes before harvesting by trypsinization. Cells were washed with PBS, and fixed with 4% paraformaldehyde (Electron Microscopy Sciences, 1571s) in PBS for 15 min at room temperature. Then 1% BSA-PBS was added and the cells were centrifuged. Fixed cells were permeabilized with 0.5% Triton X-100 in 1% BSA-PBS at room temperature for 15 min, then centrifuged. Cells were processed for EdU conjugation with 1 μM AF647-azide (Life Technologies, A10277) in 100 mM ascorbic Acid, 1 mM CuSO4, and PBS for 30 minutes at room temperature in the dark. Lastly, cells were washed and incubated in1 μg/mL DAPI (Life Technologies, D1306) overnight at 4°C. Samples were run on an Attune NxT (Beckman Coulter) and analyzed with FCS Express 7 (De Novo Software).

### P14 CD8+ T Cell Proliferation and Activation

P14 CD8+ T cell proliferation and activation assay was performed as described previously^44^. Briefly, CD8+ T cells from a P14 transgenic mouse were first isolated with CD8a microbeads according to manufacturer’s instruction. Isolated CD8+ T cells were resuspended in 1□mL 1640 medium and either labeled with 1□mL of the 10□μM Carboxyfluorescein diacetate succinimidyl ester (CFSE) for 8□min at RT for proliferation assay or kept unlabeled for activation assay. METTL3 KO, G9a KO, and control Raw 264.7 cells were seeded in 6-well plates with indicated treatments followed by 50□μg/mL Mitomycin C treatment for 30□min at 37□°C. Cells were washed with 1□mL PBS twice and then labeled with 0.4□mL of the 30□μg/mL GP33-41 peptide in PBS for 30□min at 37□°C. One hundred thousand P14 T cells were co-cultured with 2□×□10^5^ control, METTL3 KO, and G9a KO cells in 96-well plates in RPMI medium containing 50□U/mL mIL-2. The proliferation and activation of CD8+ T cells were assessed by flow-cytometry at day six.

### LC-MS/MS analysis

For ChaC pull-down samples, desalted peptide mixtures were dissolved in 30 μl 0.1% formic acid (Thermo Fisher). Peptide concentration was measured with Pierce™ Quantitative Colorimetric Peptide Assay (Thermo Fisher). In the Easy nanoLC-Q Exactive HFX setup, peptides were loaded on to a 15 cm C18 RP column (15 cm × 75 μm ID, C18, 2 μm, Acclaim Pepmap RSLC, Thermo Fisher) and eluted with a gradient of 2-30% buffer B at a constant flow rate of 300 nl/min for 30 min followed by 30% to 45% B in 5 min and 100% B for 10 min. The Q-Exactive HFX was also operated in the positive-ion mode but with a data-dependent top 20 method. Survey scans were acquired at a resolution of 60,000 at m/z 200. Up to the top 20 most abundant isotope patterns with charge ≥ 2 from the survey scan were selected with an isolation window of 1.5 m/z and fragmented by HCD with normalized collision energies of 27. The maximum ion injection time for the survey scan and the MS/MS scans was 100 ms, and the ion target values were set to 3e6 and 1e5, respectively. Selected sequenced ions were dynamically excluded for 30 seconds.

For global protein profiling, 0.5 μg of each fraction was analyzed on a Q-Exactive HF-X coupled with an Easy nanoLC 1200 (Thermo Fisher Scientific, San Jose, CA). Peptides were loaded on to a nanoEase MZ HSS T3 Column (100Å, 1.8 μm, 75 μm x 150 mm, Waters). Analytical separation of all peptides was achieved with 100-min gradient. A linear gradient of 5 to 10% buffer B over 5 min, 10% to 31% buffer B over 70 min and 31% to 75% buffer B over 15 minutes was executed at a 300 nl/min flow rate followed a ramp to 100%B in 1 min and 9-min wash with 100%B, where buffer A was aqueous 0.1% formic acid, and buffer B was 80% acetonitrile and 0.1% formic acid.

Peptides were separated with 45-min gradient, a linear gradient of 5 to 30%B over 29 min, 30 to 45%B over 6 min followed a ramp to 100%B in 1 min and 9-min wash with 100%B. LC-MS experiments were also performed in a data-dependent mode with full MS (externally calibrated to a mass accuracy of <5 ppm and a resolution of 120,000 for TMT-labeled samples or 60,000 for secretome samples at *m/z* 200) followed by high energy collision-activated dissociation-MS/MS with a resolution of 45,000 for TMT-labeled global samples and 15,000 for secretome samples at *m/z* 200. High energy collision-activated dissociation-MS/MS was used to dissociate peptides at a normalized collision energy of 32 eV (for TMT-labeled sample) or 27 eV in the presence of nitrogen bath gas atoms. Dynamic exclusion was 45 or 20 seconds. Each fraction was subjected to three technical replicate LC-MS analyses. There were two biological replicates of samples and two technical replicates were executed for each sample.

### Proteomics Data Processing and Analysis

Mass spectra were processed, and peptide identification was performed using the MaxQuant software version 1.6.10.43 (Max Planck Institute, Germany). All protein database searches were performed against the UniProt human protein sequence database (UP000005640). A false discovery rate (FDR) for both peptide-spectrum match (PSM) and protein assignment was set at 1%. Search parameters included up to two missed cleavages at Lys/Arg on the sequence, oxidation of methionine, and protein N-terminal acetylation as a dynamic modification. Carbamidomethylation of cysteine residues was considered as a static modification. Peptide identifications are reported by filtering of reverse and contaminant entries and assigning to their leading razor protein. The TMT reporter intensity found in MaxQuant was for quantitation. Data processing and statistical analysis were performed on Perseus (Version 1.6.0.7). Protein quantitation was performed using TMT reporter intensity found in MaxQuant and a one-sample t-test statistics on three technical replicates was used with a p-value of 5% to report statistically significant protein abundance fold-changes. Label-free quantification (LFQ) was for ChaC interactome and secretome data analysis.

### Analysis of Functional Category and Networks

The biological processes and molecular functions of the G9a-interacting proteins were categorized by IPA (http://www.ingenuity.com/), DAVID (http://david.abcc.ncifcrf.gov/), and STRING (http://string-db.org/).

### m^6^A RNA Immunoprecipitation Sequencing (MeRIP-Seq) and Data Analysis

m^6^A-RIP-Seq was performed as described previously with slight modifications^71^. Messenger RNA from 10 ug total RNA extracted from Ctrl, METTL3 KO and G9a KO cell samples was purified with Dynabeads Oligo (dT)_25_ (Thermo Fisher; 61006). Ten percent of 150 ng mRNA was used as Input mRNA, and the remainder was incubated with 3 ug anti-m^6^A polyclonal antibody (Synaptic Systems; 202003) which was preconjugated to Dynabeads Protein A (Thermo Fisher; 10001D) in 500uL IP buffer (50 mM Tris, pH 7.4, 150 mM NaCl, 0.1% Igepal CA-630) for 2 hours at 4°C. After washing twice with IP-buffer and twice with High-Saltwash buffer (50 mM Tris pH 7.4, 500 mM NaCl, 0.1% Igepal CA-630) for 5 minutes each, the m6A mRNA was eluted with 100 uL IP-buffer containing 6.7 mM N^6^-Methyladenosine (Sigma-Aldrich; M2780) and 40 U RNase Inhibitor (NEB, M0314S) and then recovered with RNA Clean and Concentrator-5 spin columns (Zymo; R1015).

The Input mRNA and m^6^A-IPed mRNA were subjected to library generation using the SMART-seq protocol as described (Full-length RNA-seq from single cells using Smart-seq2. Picelli et al., 2014). For the synthesis of the first strand cDNA, the mRNA was mixed with 0.25 μL RNase inhibitor and 1 μL CDS primer (5’-AAGCAGTGGTATCAACGCAGAGTACT30VN-3’) and heated to 70 °C for 2 min. Then the mixture containing 0.5 μL of 100 mM DTT, 0.3 μL of 200 mM MgCl_2_, 1 μL of 10 mM dNTPs, 0.25 μL RNase inhibitor, 1 μL of 10 μM TSO primer (5’-AAGCAGTGGTATCAACGCAGAGTACATrGrGrG-3’), 2 μL of 5X SMARTScribe RT buffer and 0.5 μL SMARTScribe reverse transcriptase (Takara, 639536) was added for performing reverse transcription. The cDNA was then amplified by Advantage Polymerase Mix (TAKARA, 639201) with IS primer (5’-AAGCAGTGGTATCAACGCAGAGT-3’). After purification with 0.8X AMPure XP beads (Fisher Scientific, A63880), the fragmentation of 100 pg cDNA was performed with EZ Tn5 Transposase (Lucigen, TNP92110). Fragments of cDNA were amplified by KAPA HiFi hotstart readymix (EMSCO/FISHER, KK2601) with the Nextera i7 primer and Nextera i5 primer. The DNA was purified with 0.8X AMPure XP beads and quantified by qPCR with KAPA Library Quantification Kit (Fisher, NC0078468). The DNA from different samples was pooled at equal molar amounts, and the final sequencing library was loaded at concentrations of 2.7 pM, and sequenced on a NextSeq 550 (Illumina) for single-read sequencing.

The raw sequencing data were demultiplexed with bcl2fastq2 v2.17.1.14 (Illumina) and the adapter was trimmed by Trimmomatic-0.32 software (Trimmomatic: a flexible trimmer for Illumina sequence data. Bolger et al., 2014). Then the Input and m6A-IP reads were mapped to human genome version hg38 by STAR v.2.5.2a (STAR: ultrafast universal RNA-seq aligner; Dobin et al., 2012), and only uniquely mapping reads at the exon level for each gene were quantified and summarized to gene counts, which were further analyzed in R v.3.6.2. After normalization, sorting, and indexing with Samtools-1.1 software^72^, the corresponding BAM files for each sample were loaded to IGV software to generate the peaks plots.

### AACT pulse-labeling and Measurements of Protein Turnover Rates

G9a KO and control Raw 264.7 cells were cultured in AACT/SILAC DMEM medium supplemented with regular lysine and arginine (K0R0) and 10% dialyzed fetal bovine serum (Thermo Fisher), 1% penicillin, and streptomycin for five passages. G9a KO, control and (1 μM) UNC0642 treated control cells were either untreated (N) or treated with low dose (100 ng/ml) LPS to induce endotoxin tolerance. The cells were washed with PBS twice to remove light medium (K0R0), and then switched to heavy AACT DMEM medium containing stable isotope-enriched D_4_-lysine and ^13^C_6_-arginine to label newly synthesized proteins. The cells were harvested at 2h, 4h, 8h, 24h, 48h, and 72h, and lysed in 8 M urea containing 50 mM Tris-HCl pH 8.0. One hundred micrograms protein from each condition was digested with trypsin, desalted, and fractionated with C18 material (High pH) into eight fractions followed by LC-MS/MS analysis.

### Supplementary Methods of AACT Pulse-labeling Translatome Analysis

Detailed description of kinetic model used for curve fitting and applied methods for data normalization

### Underlying Mathematical Model for Protein Kinetics and interpretation of Offset (*γ*)

Mathematical equations used to model synthesis and degradation of proteins in cell-lines/conditions with different doubling times were inspired from mathematical models described previously ^73–76^ and adapted to yield interpretable analytical solutions for our system. Briefly, we assumed that there is a source of amino acids and a pool of degraded waste products, both of which contain medium and heavy amino acids. Assuming we switched from medium to heavy label (i.e., *M* → *H* medium switch at *t* = 0), the heavy source pool was assumed to be the inexhaustible medium in which the cells were grown (*t* = 0). Additionally, there is a contamination of the heavy amino acid source pool via recycling so that fraction *γ*(*t*) of the amino acids in source pool is M-labelled (***Supplementary Methods* Fig. 1**). The following items are other assumptions of the model include:

a. <u>Proteins are synthesized at a constant rate (*S*)</u> using medium and heavy amino acids from the source pool. Therefore, synthesis is a zero-order process with respect to protein concentration
b. <u>Proteins are degraded at constant rate (*k*)</u> into the degradation pool. Probability of a protein being degraded is the same for old and newly synthesized protein molecules and remains constant during the lifetime of these proteins
c. <u>Rate of cell division (*k_div_*) is constant (for each cell-line/condition) and contributes to overall degradation and synthesis rates</u> (i.e., a protein with degradation rate of 0 will still show a loss of pre-existing M label because of its distribution equally to daughter cells. Similar thing occurs for newly synthesized proteins as intensity of H-label will increase owing to cell doubling even if protein’s per cell concentration is constant)
d. <u>Cells are at steady state.</u> The concentration per cell of any given protein remains constant during the experiment. Therefore, synthesis and degradation rates of a protein are equal (i.e., *S* = *k* + *k_div_*) and first order labelling kinetics were adopted for curve fitting and intensities were fitted to exponential equations.

**Figure 1:**
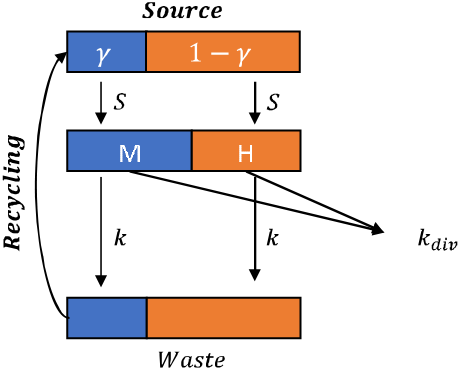
Schematic representation of the synthesis and degradation model taking amino-acid recycling as well as effect of cell division into account

**Figure 2:**
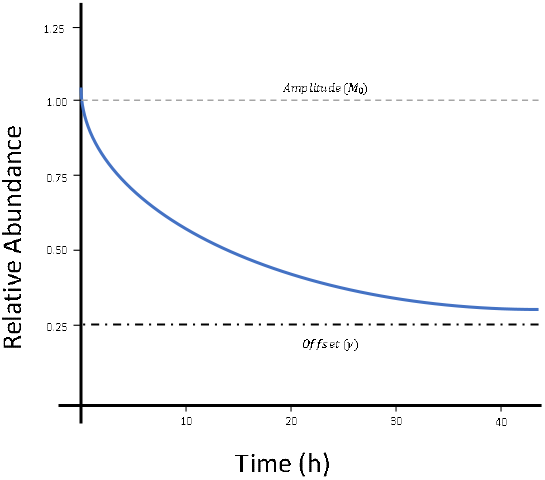
Illustration of the exponential decay with an asymptotic offset, *M*(*t*) = (*M*_0_ – *γ*)*e*^−(*k*+*k_div_*)*t*^ + *γ*. The amplitude (*M*_0_ = 1) and offset (*γ* = 0.25) are shown by the horizontal dotted lines.

**Figure 3:**
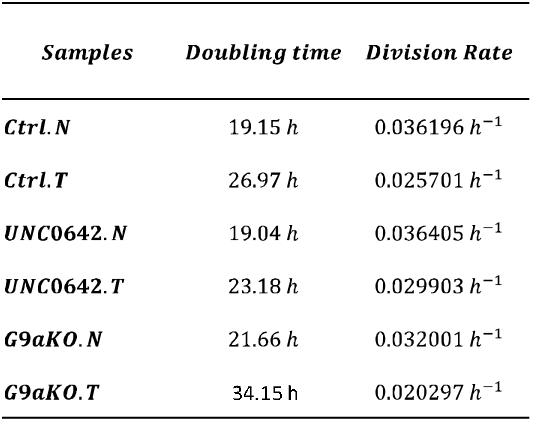
Cell division rates (*k_div_*) for the six conditions were determined using CCK-8 kit.

All model equations/diagrams are for a protein *i* with the index dropped for clarity. Master equations describing the system are as follows:

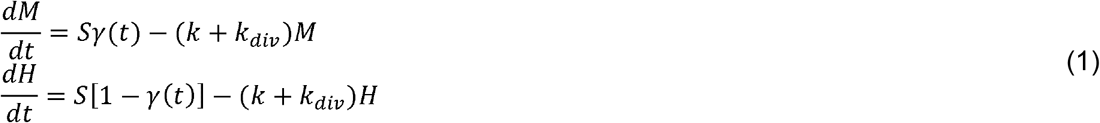

where *M* = *M*(*t*) and *H* = *H*(*t*) are dimensionless abundances of medium and heavy proteins, respectively.

### Constant Offset (*γ*)

In steady state we expect the contamination fraction to stabilize at a certain level, *γ*(*t*) = *γ* = *const*. In such case the equations (1) have a trivial solution:

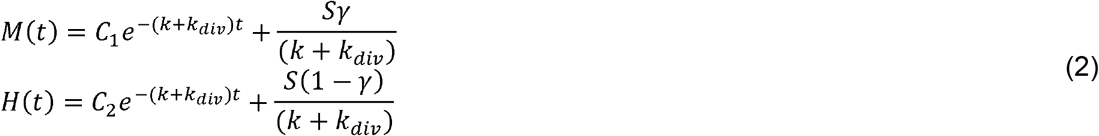

We require a stationary solution where the total amount of protein is constant, *H*(*t*) + *M*(*t*) = *const*.

Hence, 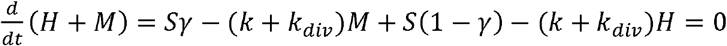 and we find *S* = (*k* + *k_div_*)(*H* + *M*). If we normalize our data, such that *H*(*t*) + *M*(*t*) = 1 we find that:

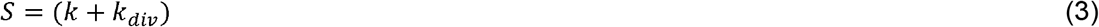

At time *t* = 0 we have *H*(*t*) = *H_o_* and *M*(*t*) = *M_o_*. Therefore, using equations (2) and (3), we can find *C*_1_ and *C*_2_ and plugging their value back into equations in (2), we get:

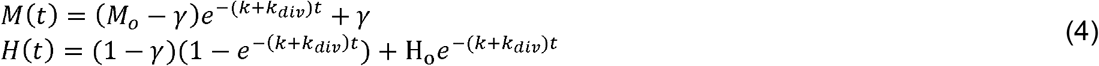

Where *M_o_* and *H_o_* are intensities of medium and heavy labeled proteins at *t* = 0 and should ideally be 1 and 0 respectively (assuming *M* → *H* label switch at *t* = 0). However, they have been left as free variables in the equations above to account for offset (*γ*) caused by amino acid recycling, experimental variation, and errors in data fitting. The degradation curve *M*(*t*) follows a simple exponential decay with an offset (*γ*) which corresponds to the (constant) fraction of contamination of the amino acid source pool **(*Supplementary Methods Fig. 2*)**.

Analytical solution for determining protein half-life from equations in (4) is as follows:

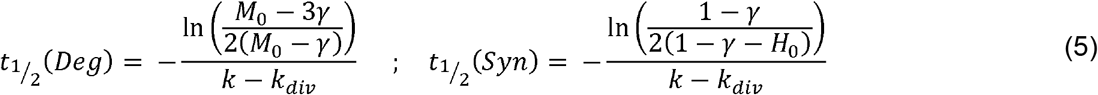

However, because of errors while fitting free variables (*γ*, *k* and *H*_0_ or *M_o_*), the solutions are not stable.

Therefore, we require a numerical solution to determine protein half-life instead. Numerical solutions show that in the initial part of the decay, *γ*(*t*) is small and *S* ≈ *k* + *k_div_*. Hence, we can rewrite equations in (4):

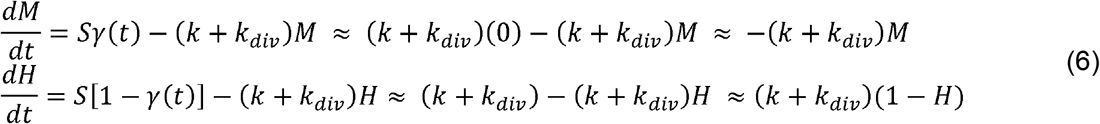

These are trivial equations of pure degradation and synthesis with exponential solutions of the form (at *t* = 0) as shown:

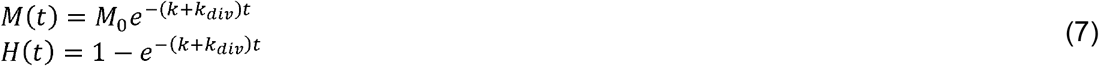

From this we can find an approximate relation between half-life and the degradation/synthesis coefficient given by:

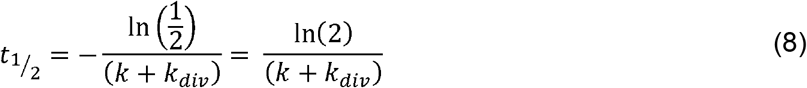

### Asymptotic Offset (*γ*)

The constant *γ* case can be generalized to a scenario in which *γ*(*t*) increases from the initial *γ*(0) = 0 until the entire system reaches an asymptotic equilibrium. This was consistent with our data where observed *M*(*t*) and *H*(*t*) reach an equilibrium for most of the proteins. In this asymptotic offset (*γ*) state we have

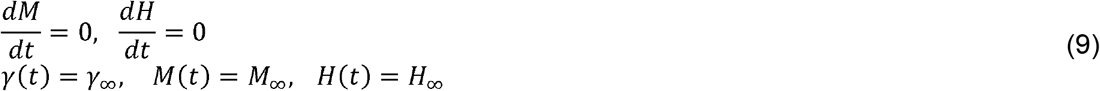

Substituting these into the original equations (1) yields

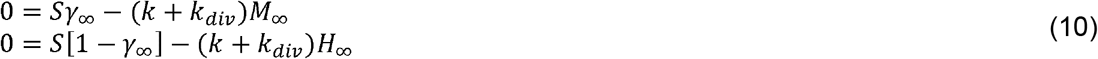

Which gives *M*_∞_ + *H*_∞_ = *S*/(*k* + *k_div_*). Again, we require a stationary solution with normalization *M*(*t*) + *H*(*t*) = 1, hence *S* = (*k* + *k_div_*). We reach a similar interpretation of the offset: it is explained by the asymptotic contamination fraction, *M*_∞_ = *γ*_∞_.

### Exponential offset (*γ*)

One possible functional form for *γ*(*t*) is an exponential growth, *γ*(*t*) = *γ*_∞_(1 – *e^−kt^*). In such case the first equation of (1) assumes the form

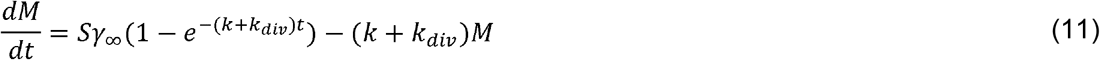

which can be solved analytically

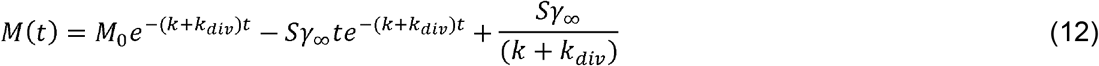

and in a case of balanced synthesis and degradation (*S* = (*k* + *k_div_*)) we obtain

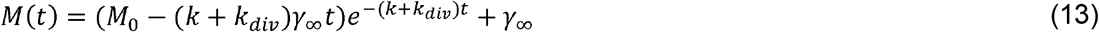

This is a modified exponential decay with a constant asymptotic offset, *γ*_∞_. The shape of this curve resembles a simple exponential decay with an offset. The offset, *γ*_∞_, is interpreted again as the asymptotic contamination fraction of the source pool.

### AACT/SILAC pulse-labeling data extraction and normalization

Cell lysates were collected at 2h, 4h, 8h, 24h, 48h and 72h from wildtype (Ctrl), G9a knockout (G9a-KO) and 1 *μM* UNC0642-treated Raw 264.7 cells under N and T conditions, digested with trypsin, fractionated by HPLC and finally subjected to LC-MS/MS analysis. To correct for errors introduced during sample-preparation/data-acquisition, protein intensities were normalized based on the premise that the sum of K0/R0 (M) and the K4/R6 (H) intensities should be constant across different time-points. After normalization, curves for estimation of turnover rates, determined from the kinetics of AACT label incorporation or loss, were fitted based on the assumption of exponential protein degradation (or synthesis) and constant offset (*γ*).

Proteins identified in at-least 4 out of 6 time points were selected for curve fitting using the model described above. The turnover rates (*k*_deg_/*k_syn_*), curve maxima (*M_o_*/*H_o_*), offsets (*γ*) and goodness-of-fit statistics (*SSE, RMSE, rsquared, adjrsquared*) were obtained for each protein using a nonlinear least square (NLS) method in MATLAB (vR2017b). To remove poor quality quantitative data, different filter criteria for *k*, *R*^2^, *M_o_*/*H_o_* and *γ* were applied, and curves that were at the border of passing these filter criteria were manually inspected. The final filtering criteria were based on the goal to remove spectra that showed a high variation of data points along the fitted curve (*M_o_*/*H_o_* and *R*^2^), a high offset (*γ*), or resulted in turnover rates (*K*) that simply could not be determined accurately considering the pulse time-points we have chosen in the experimental design. Eventually, only entries were kept that met following filter criteria: *k_deg_*/*k_syn_*: [0,5]; *M_o_*: [0.65, 1.5]; *H_o_*: [-0.5, 0.50]; *γ*: [-0.2, 0.5] and coefficient of determination *R*^2^ ≥ 0.8. Subsequently, protein synthesis 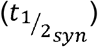 and degradation 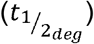 half-lives were estimated using equation (8). Finally, synthesis and degradation half-lives were combined to obtain median protein turnover time/half-life 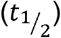.

## EXTENDED DATA FILES

**Extended Data Fig. 1. ET-phenotypic G9a interactome consist of SARS-CoV-2-upregulated proteins**. **a**. Certain known components of G9a complex are overexpressed in COVID-19 patients with high viral load. Heatmap depicts mRNA expression of core G9a complex members that undergo significant (FDR corrected P < 0.01) transcriptional regulation in response to SARS-CoV-2 infection. COVID-19 patient samples are stratified into ‘none’, ‘low’, ‘medium’, and ‘high’ based on viral load as defined by presence of SARS-CoV-2 RNA (detected using NGS & RT-PCR from nasal swabs). ‘Other viral infection’ category includes patients with unrelated viral infections, which serves as an additional control. Violin plots on the right show median, quartiles and minima and maxima bounds (TPM, transcripts per million, shown on y axis) for various G9a complex members in Covid-19 patients in context of viral load [adapted using data from ref. ^15^]. **b**. SARS-CoV-2 upregulated pathways and associated G9a interactors that were identified by UNC0965 ChaC-MS in ET. **c**. Depletion of either METTL3 or G9a caused similar reduced ET and resensitized the ET macrophages to LPS stimulation

**Extended Data Fig. 2. G9a and METTL3 co-upregulate translation of m6A-marked mRNA subsets**. **a**. Integrative genomics view (IGV) plots of m^6^A peaks at CD274 (PD-L1) mRNA in different inflammatory conditions (N, NL, or TL). The y axis shows sequence read number (blue peaks represent the intensity of m^6^A) in either ‘input’ or m^6^A immunoprecipitates ‘IP’. Two biological replicates are included as ‘input1 or IP1 or input2 or IP2’. Blue boxes are exons, and blue lines are introns. **b**. Enriched functional category/pathways overrepresented by G9a/METTL3-co-upregulated m^6^A mRNA and proteins in ET. **c**. qPCR measurements of mRNA of G9a/METTL3-co-downregulated m^6^A-tagged genes RGS13, CD274, BIRC5, and CX3CR1; mRNA expression was normalized to GAPDH. **d**. G9a affects the total m^6^A level in macrophage and monocyte cells. (Left): Knocking down G9a (shRNA) in Raw 264.7 macrophage cells reduced the total m^6^A level of mRNA in endotoxin tolerance condition. *p<0.05. (Right): Knocking out G9a or METTL3 in THP-1 monocyte cells reduced the total m^6^A level of mRNA in endotoxin tolerance condition. *p<0.05, **p<0.01. **e**. Quantitative proteomic profiling of G9a/METTL3-co-regulated proteins. (Left) Principal component assay to present the samples separations. (middle and right) volcano plots to present specific METTL3 or G9a interactors (purple) in endotoxin tolerance (ET or TL) compared with control sample. The log_2_Fold changes >1 or <-1, and −log10 p-value > 2 (p<0.01) were the threshold applied in the data processing. Each condition has two biological and three technical replicates.

**Extended Data Fig. 3**. **a**. Constitutively active G9a stabilized Myc protein. (upper) Myc protein expression was diminished in G9a KD cells under TL conditions; actin was the loading control. Tolerized Raw 264.7 cells were treated with 1 μM CHX for indicated time.(lower) c-Myc accumulation was investigated by applying a timecourse MG-132 (10 μM) treatment on LPS-tolerant cells. b. METTL3 is post-translationally methylated and phosphorylated at multiple sites. b. HA-G9a and Flag-METTL3 were overexpressed in 293T cells and purified by Flag IP. Cell lysates and IP eluates were subjected to immunoblotting with antibodies that recognized pan mono-and di- or tri-methylated lysine. METTL3 was methylated. c. LC-MS/MS of METTL3 protein in the Flag IP product. Flag-METTL3 IP sample was further fractionated on 8% SDS-PAGE gel, and the protein band at 70 kDa was excised, in-gel digested with trypsin, and analyzed by LC-MS/MS. Red indicates covered METTL3 sequence; modified lysine or serine are in green and underlined. d. A summary of identified methylation (four sites) and phosphorylation sites (two sites) of METTL3 protein. MS raw files were subjected to database search (Maxquant software) with oxidation (O), protein N-terminus acetylation, acetyl (K), methyl(KR), dimethyl (K), tri-methyl (K) as variable modifications. A separate search with only oxidation(O), protein N-term acetylation and phosphorylation (STY) was performed simultaneously. FDR for peptide identification is 0.01, modified sites with the probabilities score >0.85 were considered. Peptide mass to charge value (m/z), charge, and score were also listed.

**Extended Data Fig. 4. Reproducibility of protein turnover rate determination by AACT based pulse-labeling strategy and identification of G9a translated proteins and pathways a**. Correlation matrix depicts color-coded Pearson’s correlation coefficients for protein turnover time [half-life, hrs] determined from synthesis and degradation curves for N and T conditions. The boxplots (10th–90th percentile) show the covariance for protein half-lives across synthesis and degradation curve pairs within a sample, N/T conditions and cell culture replicates. **b**. Examples of the reproducibility of turnover determination across cell culture replicates are displayed; Mitogen-Activated Protein Kinase 3 (MAPK3) and Transforming Growth Factor Beta 1 (TGBβ1). Points where synthesis and degradation curves intersect depict apparent half-life which is corrected for differences in cell division rates (between the 6 conditions) to yield actual protein synthesis and degradation rates for the protein (shown in the table). **c**. Principle component analysis showing clear difference among cellular proteostasis landscapes of N and TL conditions. Genetic/pharmacologic inhibition of G9a appears to have large scale effect on protein synthesis/degradation rates compared with wild type controls. **d**. Tuckey’s plot shows pairwise differences in mean turnover time between different conditions globally. (related to **Fig. 1c**). **e**. Pairwise comparisons of protein turnover time (against *Ctrl.N*) were performed to identify 3994 proteins that were translated in G9a dependent manner in N and T condition. G9a/ET translated proteins (>2FC increase (red)/decrease (blue) in half-life) are highlighted. **f**. Results of canonical pathway analysis for genes highlighted in (**e**) are summarized. Sizes of the circles depicts number of proteins in each pathway, and color depicts log transformed p-value. G9a primarily controls turnover of proteins involved in immunity, cell cycle, translation/proteostasis, cellular energetics, RNA biogenesis and coronavirus related pathways. G9a inhibition appears to reverse the effect of ET on global proteostasis landscape. **g**. G9a may control global protein turnover by multiple pathways because several of the differentially translated proteins are G9a interactors, substrates or m^6^A regulated targets. **h**. Table showing effect of G9a inhibition on select proteins of interest.

**Extended Data Fig. 5. G9a/ET affect net protein turnover by different pathways to varying degrees**. **a**. G9a/METTL3 regulated m^6^A targets. **b**. Coronavirus related pathway proteins and SARS-CoV2 host interactors. **c**. Immune System Related Pathways. **d**. Cell Cycle Related Pathways. **e**. RNA Biogenesis Related Pathways. **f**. Cellular Energetics Related Pathways. **g**. Translation/Proteostasis Related Pathways. **h**. Miscellaneous Pathways

**Extended Data Fig. 6. PRC2 complex members are overexpressed in Covid-19 patients with high viral load. a**. Heatmap depicting mRNA expression of PRC2 complex members in response to SARS-CoV-2 infection. COVID-19 patient samples have been stratified into ‘none’, ‘low’, ‘medium’, and ‘high’ based on viral load as defined by presence of SARS-CoV-2 RNA (detected using NGS & RT-PCR from nasal swabs of patients). ‘Other viral infection’ category includes patients with unrelated viral infections and serves as an additional control. **b**. Violin plots indicating median and quartiles as well as minima and maxima bounds (TPM, transcripts per million, shown on *y* axis) for various PRC2 complex members in COVID-19 patients in context of viral load [a. and b. adapted using data from ref.^15^]. **c**. Heatmap showing median turnover time for PRC2 complex members identified in our translatome dynamics studies.

**Extended Data Fig. 7**. **a**. Genetic perturbation of several G9a associated proteins adversely affects SARS-CoV-2 replication and infection [adapted using data from ref. ^13,30^]

**Extended Data Table 1**. List of G9a interactors identified by UNC0965 ChaC-MS in the TL/ET macrophages.

**Extended Data Table 2**. List of the proteins identified in both the UNC0965-captured complex and the Flag-METTL3 immunoprecipitated complex from TL macrophages.

**Extended Data Table 3**. Curve fitting parameters, list of differentially turned over proteins, and pathway enrichment analysis of G9a translated proteins in wildtype (ctrl), G9a-KO and UNC0642 treated non-stimulated (N) and endotoxin tolerant (T/ET) macrophage cells.

**Extended Data Table 4**. COVID-19-characteristic proteins that are suppressed by G9a or Ezh2 inhibitor in ET macrophages.

