## Supplemental Figures for "Novel gene-specific translation mechanism of dysregulated, chronic inflammation reveals promising, multifaceted COVID-19 therapeutics"

### Extended Data Fig. 1.

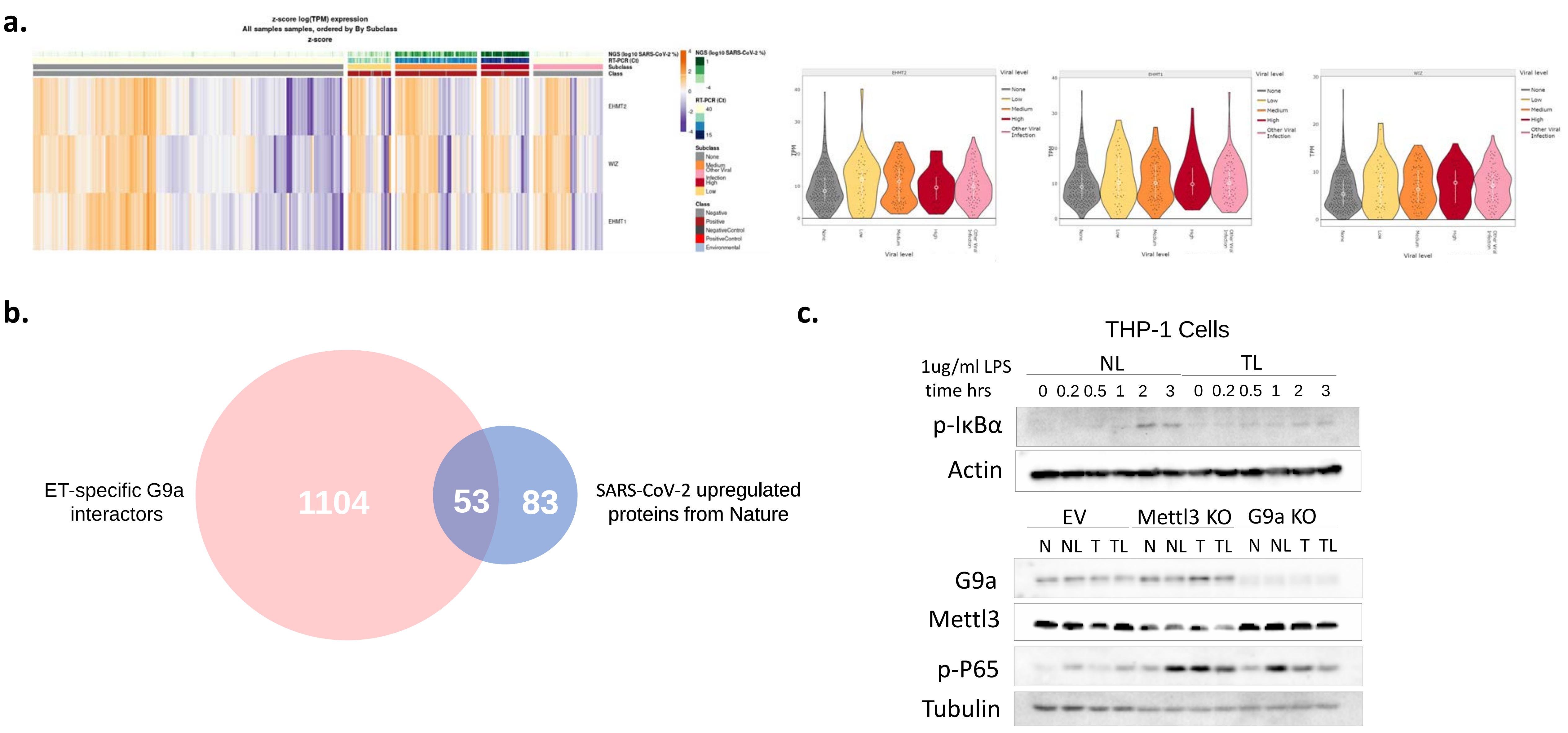

### Extended Data Figure 2

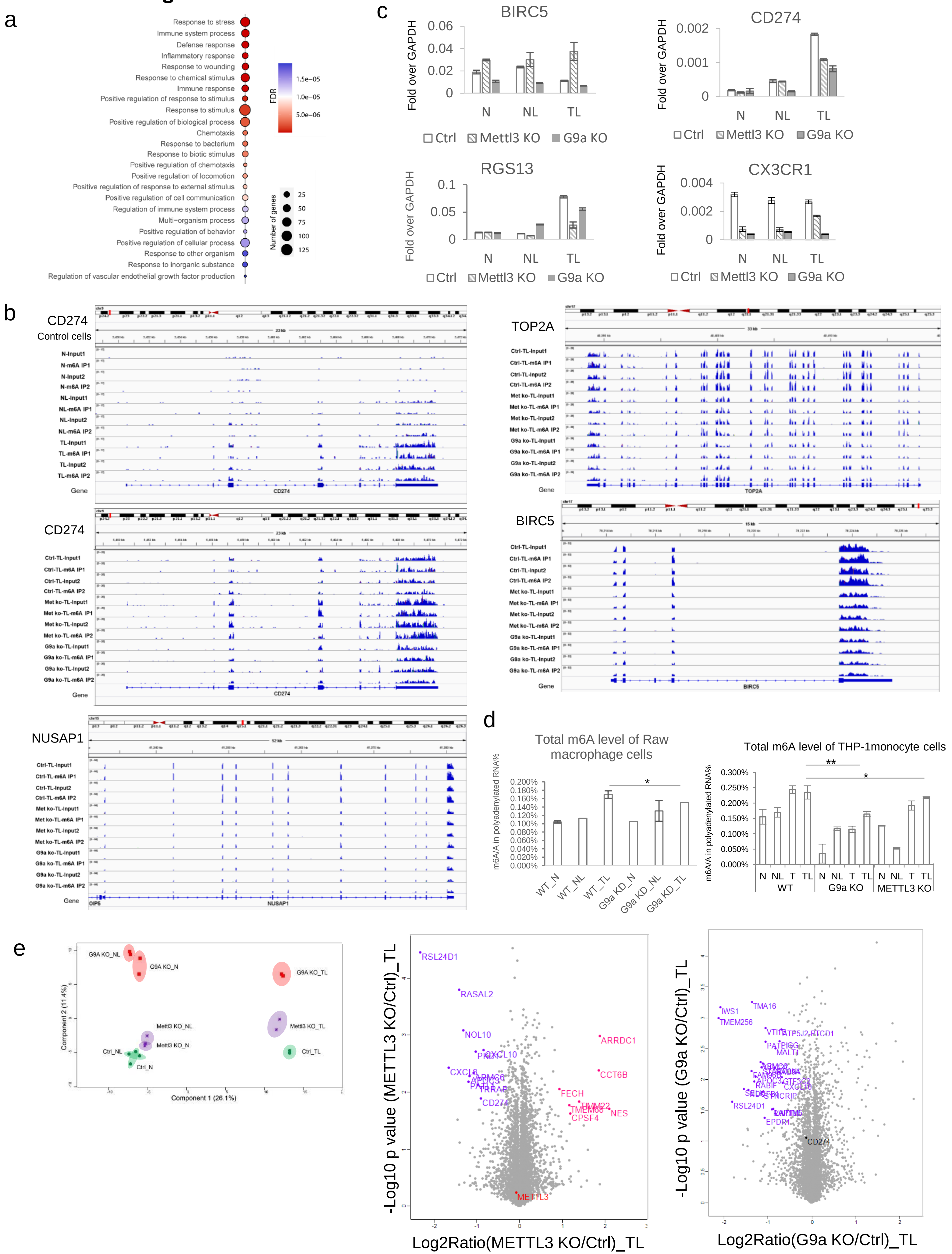

### Extended Data Figure 3

a

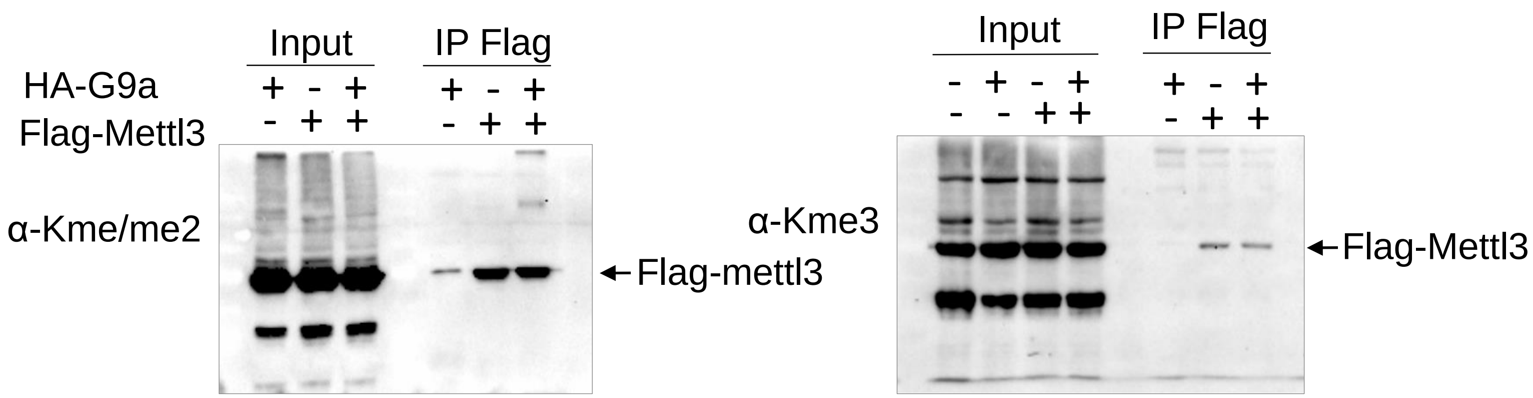


b

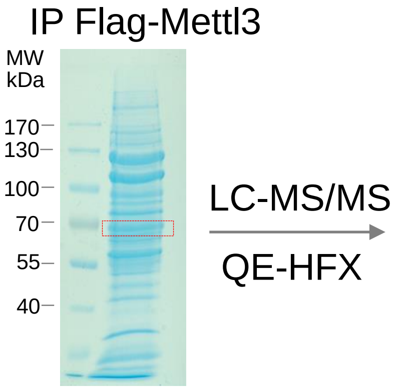


Mettl3 Sequence coverage: 89.1%

MSDTWSSIQA HKKQLDSLRE RLQRRRKQDS GHLDLRNPEA ALSPTFRSDS 50
PVPTAPTSGG PKPSTASAVP ELATDPELEK KLLHHLSDLA LTLPTDAVSI 100
CLAISTPDAP ATQDGVESLL QKFAAQELIE VKRGLLQDDA HPTLVTYADH 150
SKLSAMMGAV AEKKGPGEVA GTVTGQKRRA EQDSTTVAAF ASSLVSGLNS 200
SASEPAKEPA KKSRKHAASD VDLEIESLLN QQSTKEQQSK KVSQEILELL 250
NTTTAKEQSI VEKFRSRGRA QVQEFCDYGT KEECMKASDA DRPCRKLHFR 300
RIINKHTDES LGDCSFLNTC FHMDTCKYVH YEIDACMDSE APGSKDHTPS 350
QELALTQSVG GDSSADRLFP PQWICCDIRY LDVSILGKFA VVMADPPWDI 400
HMELPYGTLT DDEMRRLNIP VLQDDGFLFL WVTGRAMELG RECLNLWGYE 450
RVDEIIWVKT NQLQRIIRTG RTGHWLNHGK EHCLVGVKGN PQGFNQGLDC 500
DVIVAEVRST SHKPDEIYGM IERLSPGTRK IELFGRPHNV QPNWITLGNQ 550
LDGIHLLDPD VVARFKQRYP DGIISKPKNL 580

c

Modified peptides of Mettl3

| Sites | Modifications | Probabilities | m/z | Charge | Score |
|---|---|---|---|---|---|
| K215me2 | Dimethyl (K) | K(1)HAASDVDLEIESLLNQQSTK | 589.5616 | 4 | 130.89 |
| K215me | Methyl (KR) | K(1)HAASDVDLEIESLLNQQSTK | 781.0721 | 3 | 182 |
| K215me | Methyl (KR) | K(1)HAASDVDLEIESLLNQQSTK | 586.0572 | 4 | 121.47 |
| K281me | Methyl (KR) | AQVQEFCDYGTK(1) | 730.3281 | 2 | 118.18 |
| K286me | Methyl (KR) | AQVQEFCDYGTK(0.011)EECMK(0.989) | 713.6382 | 3 | 123.03 |
| K327me | Methyl (KR) | HTDESLGDCSFLNTCFHMDTCK(1) | 673.2751 | 4 | 200.99 |
| K327me | Methyl (KR) | HTDESLGDCSFLNTCFHMDTCK(1) | 897.3652 | 3 | 184.2 |
| K327me | Oxidation (M),Methyl (KR) | HTDESLGDCSFLNTCFHMDTCK(1) | 677.2751 | 4 | 182.15 |
| K327me | Oxidation (M),Methyl (KR) | HTDESLGDCSFLNTCFHMDTCK(1) | 902.6971 | 3 | 296.22 |
| pS350 | Phospho (ST) | DHT(0.127)PS(0.873)QELALTQSVGGDSSADR | 784.3433 | 3 | 202.47 |
| pS219 | Phospho (ST) | KHAAS(1)DVDLEIESLLNQQSTK | 803.0578 | 3 | 103.33 |

### Extended Data Fig. 4

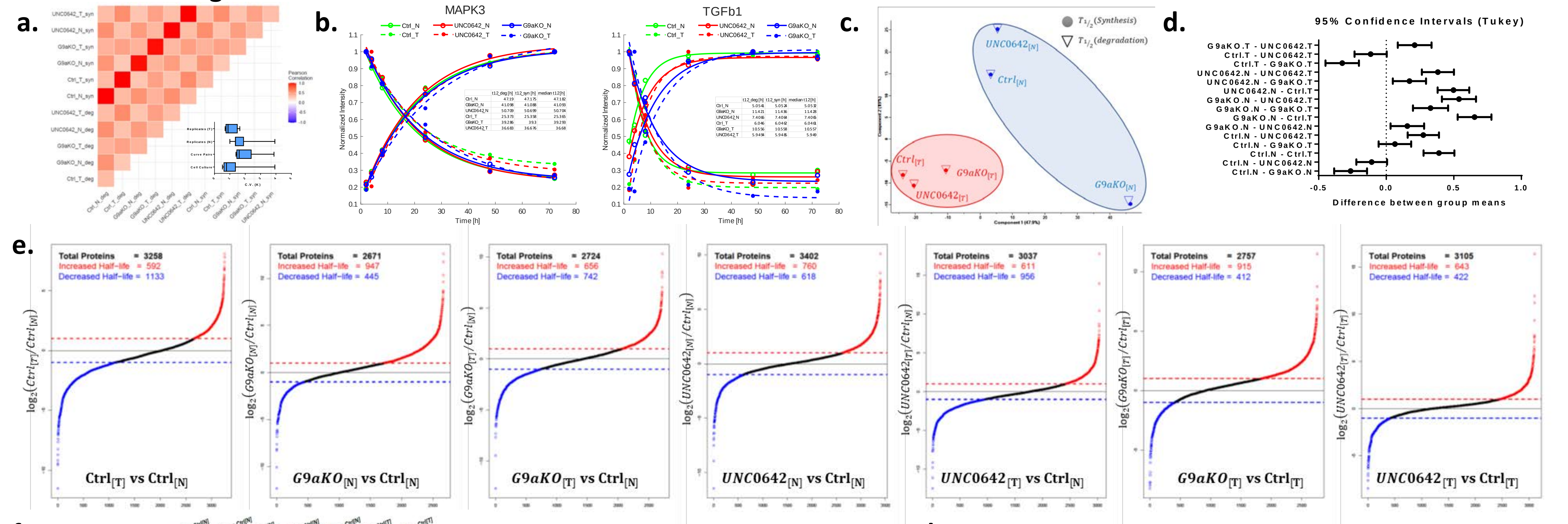


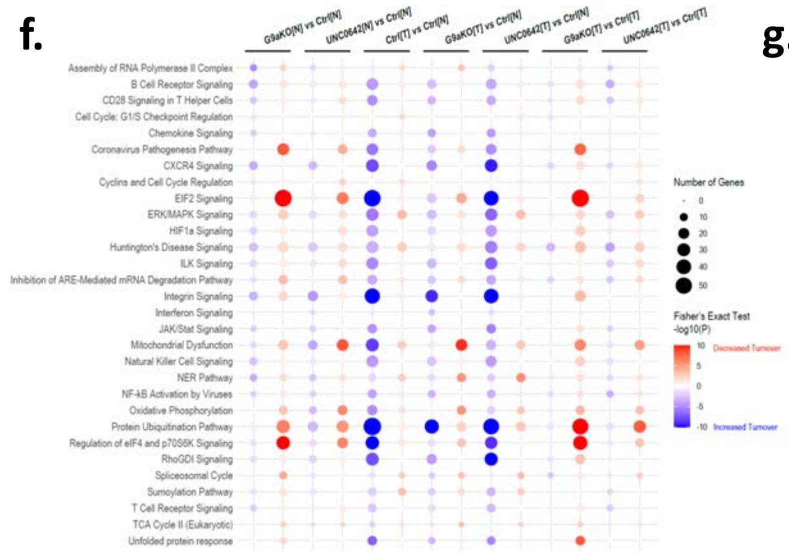


**g.**

| Annotation | | | Total Filtered Curves (6243) | G9a Translated Genes (3994) |
|---|---|---|---|---|
| m6A target (meRIP-Seq) | METTL3-KO.N vs Ctrl.N | Up | 89 | 46 |
| | | Down | 37 | 20 |
| | METTL3-KO.TL vs Ctrl.TL | Up | 76 | 48 |
| | | Down | 32 | 14 |
| | G9aKO.N vs Ctrl.N | Up | 158 | 85 |
| | | Down | 92 | 49 |
| | G9aKO.TL vs Ctrl.TL | Up | 182 | 116 |
| | | Down | 93 | 58 |
| | m6A regulated (Combined) | Coregulated | 159 | 85 |
| | | G9a Regulated | 274 | 170 |
| | | METTL3 Regulated | 39 | 27 |
| ET specific G9a/GLP Interactor (ChaC Identified) | | | 1184 | 774 |
| G9a/GLP Substrate (known/putative) | | | 976 | 760 |
| SARS-CoV2 Host Interactor | | | 216 | 157 |
| Covid19 Markers | | | 11 | 4 |

**h.**

| Category | Gene Name | Median Turnover Time [Hrs] | | | | | | Z-scored | | | | | |
|---|---|---|---|---|---|---|---|---|---|---|---|---|---|
| | | Ctrl.N | G9aKO.N | UNC0642.N | Ctrl.T | G9aKO.T | UNC0642.T | Ctrl.N | G9aKO.N | UNC0642.N | Ctrl.T | G9aKO.T | UNC0642.T |
| COVID19 Markers | ANPEP | 36.4687 | 89.4377 | 398.901 | 21.0017 | 41.0844 | 31.5726 | -0.45358047 | -0.09288282 | 2.0144382 | -0.55890454 | -0.4221494 | -0.4869209 |
| | DST | NA | NA | NA | NA | 9.95257 | NA | NA | NA | NA | NA | NaN | NA |
| | FABP4 | NA | NA | NA | 9.50274 | 5.76144 | 13.0148 | NA | NA | NA | 0.02106627 | -1.0103667 | 0.9893004 |
| | IL1RN | 56.8433 | 48.4577 | 175.306 | 9.72533 | 11.1528 | 9.27979 | 0.07881449 | -0.05208026 | 1.9279541 | -0.65667193 | -0.6343899 | -0.6636266 |
| | PLAU | 7.51876 | 5.45226 | NA | 12.4297 | 5.45335 | NA | -0.0591698 | -0.68699873 | NA | 1.43283611 | -0.6866676 | NA |
| | SPP1 | NA | 1.53846 | NA | NA | NA | 5.22894 | NA | -0.70710678 | NA | NA | NA | 0.7071068 |
| | SQSTM1 | 6.56646 | 6.77021 | 6.54385 | 6.14809 | 13.1639 | 6.49783 | -0.38468942 | -0.30994146 | -0.3929842 | -0.53817314 | 2.0356553 | -0.4098671 |
| | TNF | NA | NA | 5.27208 | NA | 7.75642 | 9.58636 | NA | NA | -1.0465553 | NA | 0.100736 | 0.9458193 |
| | CCL4 | NA | NA | NA | 0.983788 | NA | NA | NA | NA | NA | NaN | NA | NA |
| | CXCL2 | NA | NA | NA | NA | 30.976 | NA | NA | NA | NA | NA | NaN | NA |
| | IL1B | NA | NA | NA | NA | 11.9718 | NA | NA | NA | NA | NA | NaN | NA |
| Cell Cycle | ALCAM | 13.7415 | 22.348 | 16.0005 | 14.1334 | 8.34321 | 15.5351 | -0.2824917 | 1.6237074 | 0.2178399 | -0.1956923 | -1.4781247 | 0.1147614 |
| | CA2 | NA | NA | NA | NA | 158.814 | NA | NA | NA | NA | NA | NaN | NA |
| | CD300A | 6.0826 | NA | NA | 1.14781 | NA | NA | 0.7071068 | NA | NA | -0.7071068 | NA | NA |
| | SLC2A1 | 8.1689 | 15.7842 | 15.5189 | 4.6555 | 5.80882 | 7.5378 | -0.2897817 | 1.2751733 | 1.2206538 | -1.0117903 | -0.7747814 | -0.4194736 |
| | TOP2A | 9.12051 | 2.49106 | NA | NA | NA | NA | 0.7071068 | -0.7071068 | NA | NA | NA | NA |
| | NUSAP1 | 3.19051 | 4.38949 | NA | NA | NA | NA | -0.7071068 | 0.7071068 | NA | NA | NA | NA |
| Immune Response | CD274 (PD-L1) | NA | NA | NA | 8.48666 | NA | 28.414 | NA | NA | NA | -0.7071068 | NA | 0.7071068 |
| | C5AR1 | NA | NA | NA | 2.75021 | 8.17933 | 13.6267 | NA | NA | NA | -0.9994402 | -0.00111862 | 1.000559 |

### Extended Data Fig. 5

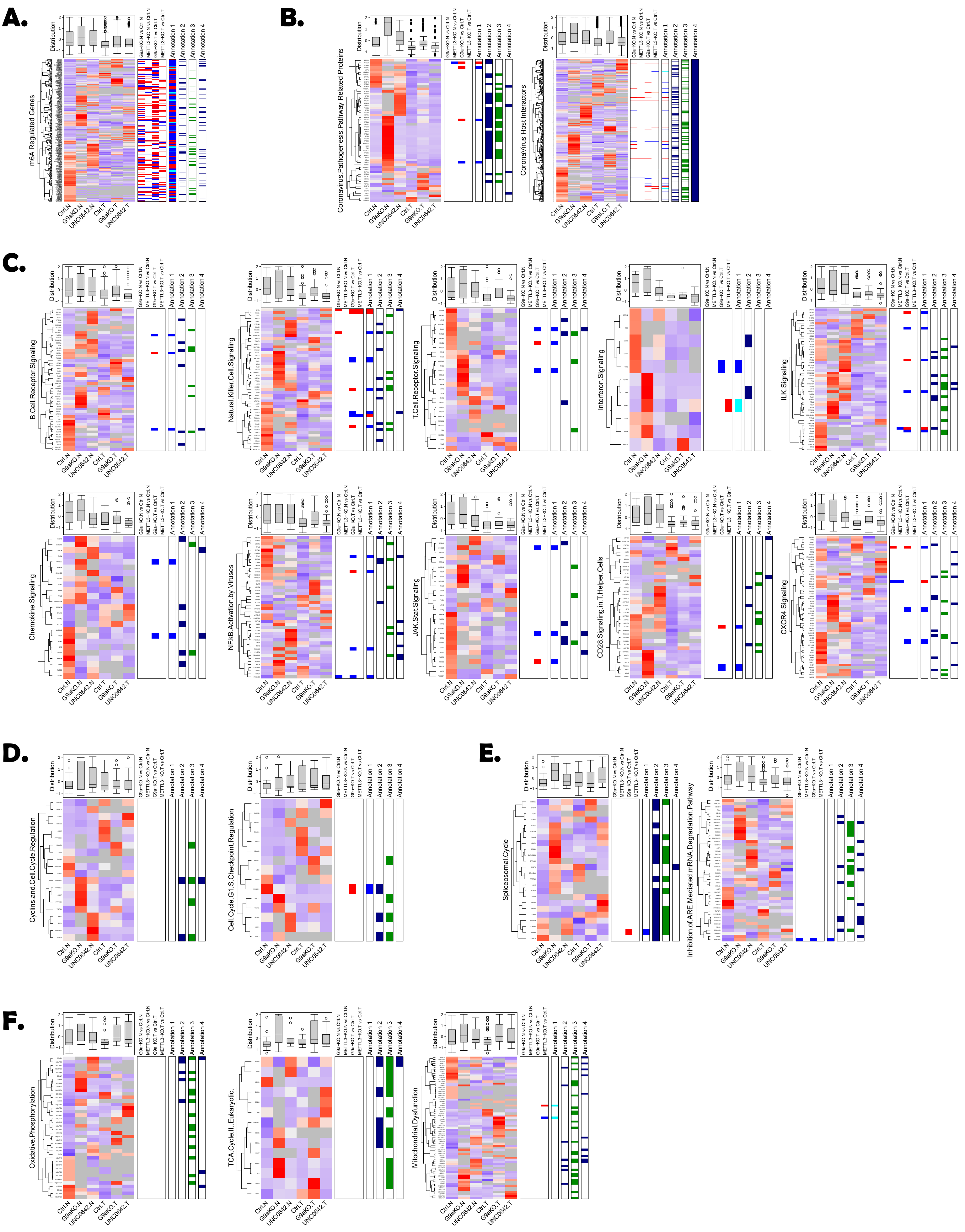

### Extended Data Fig. 5 (Cont'd)

## G.

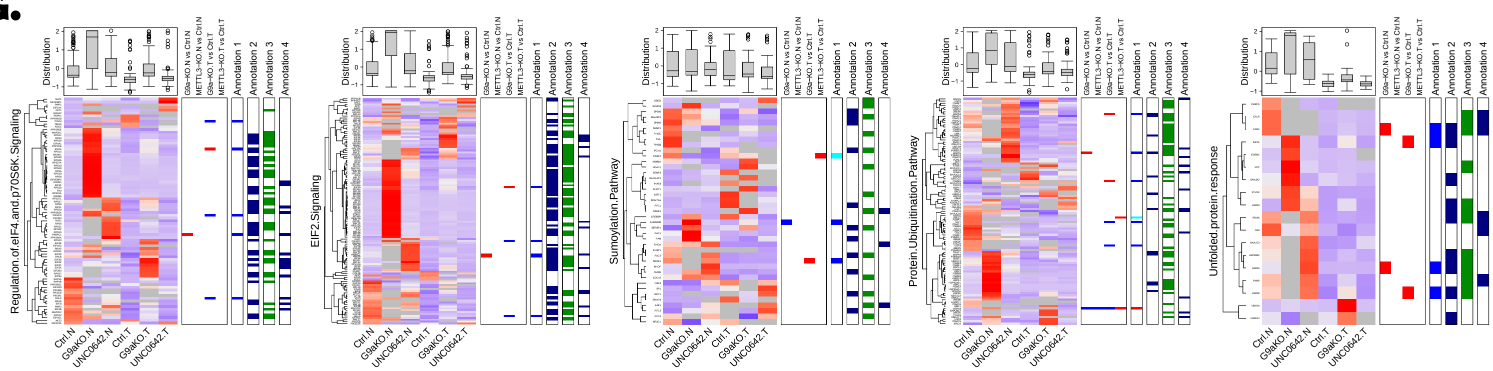


## H.

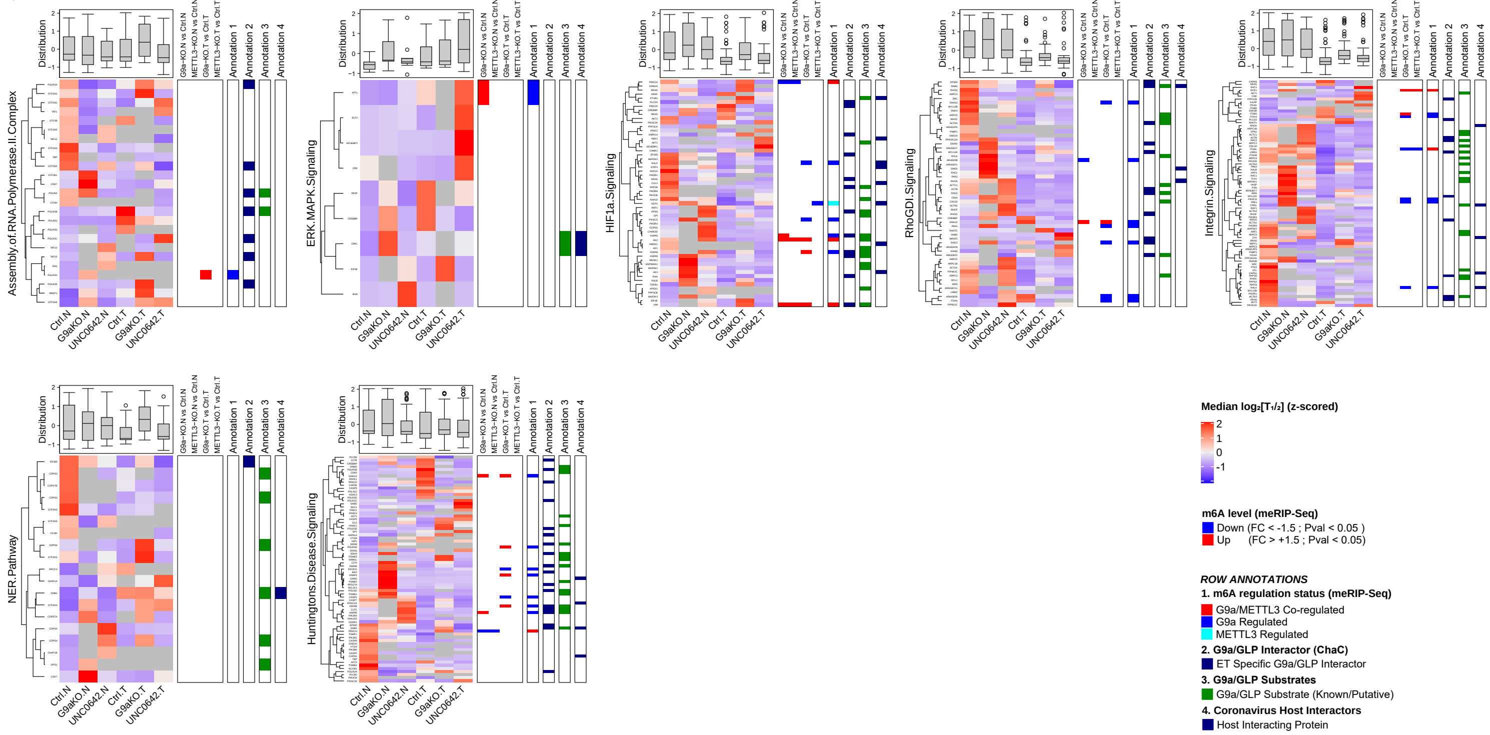

### Extended Data Fig. 6

a.

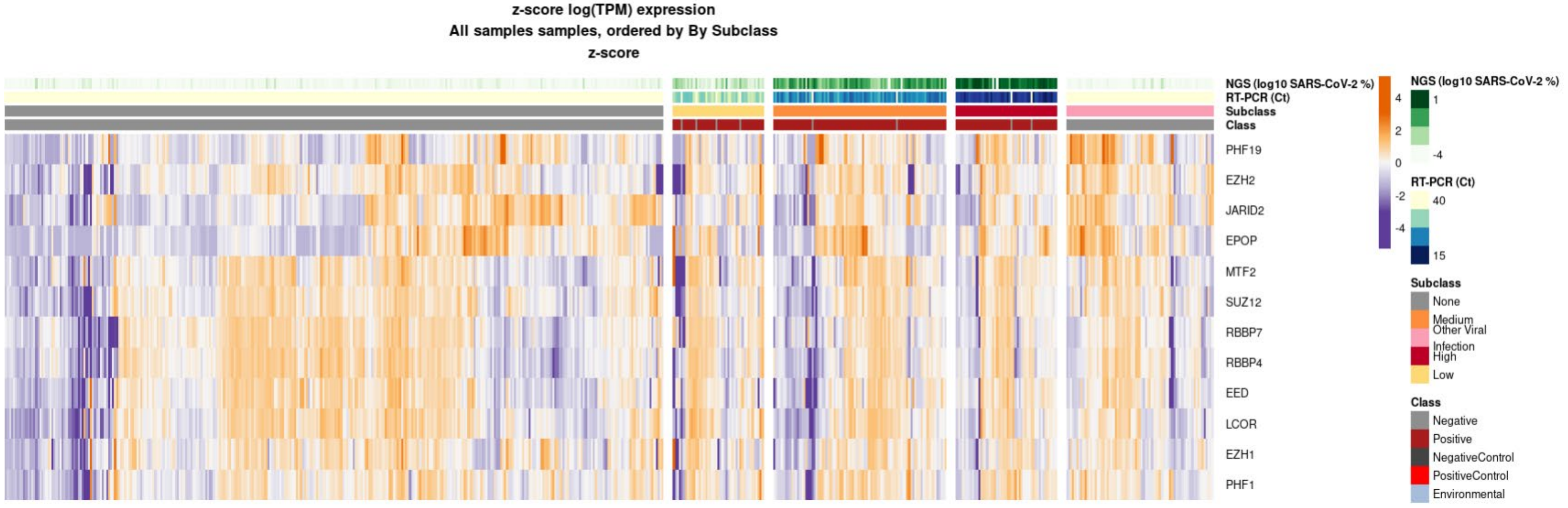


c.

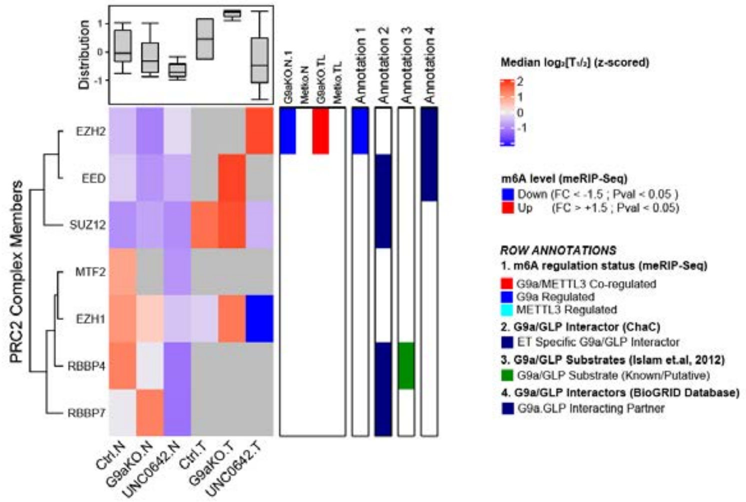


b.

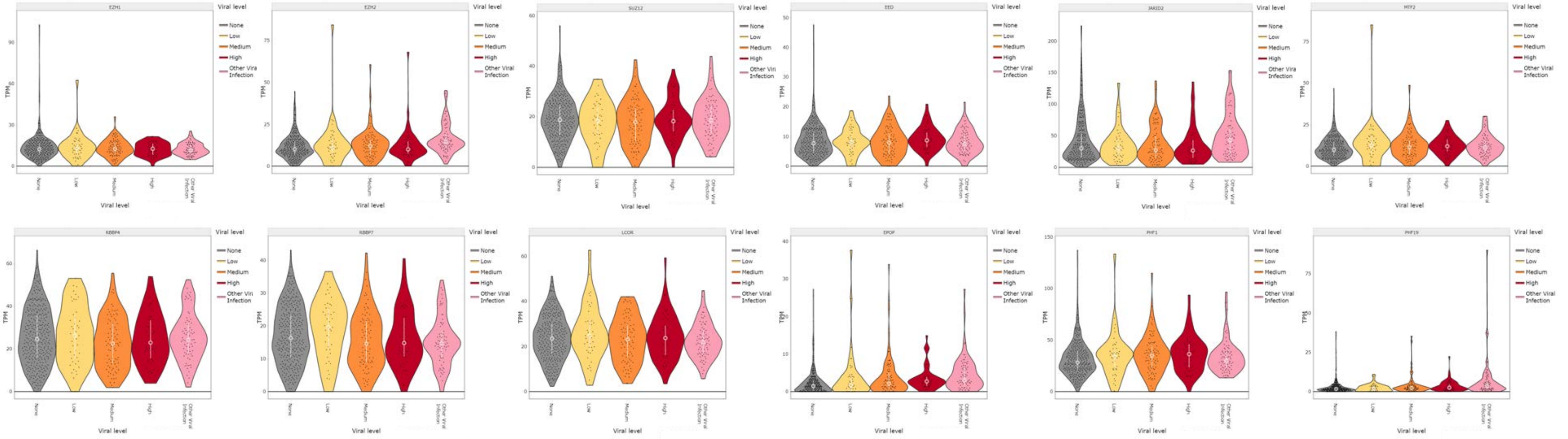

**Extended Data Fig. 7**

**a.**

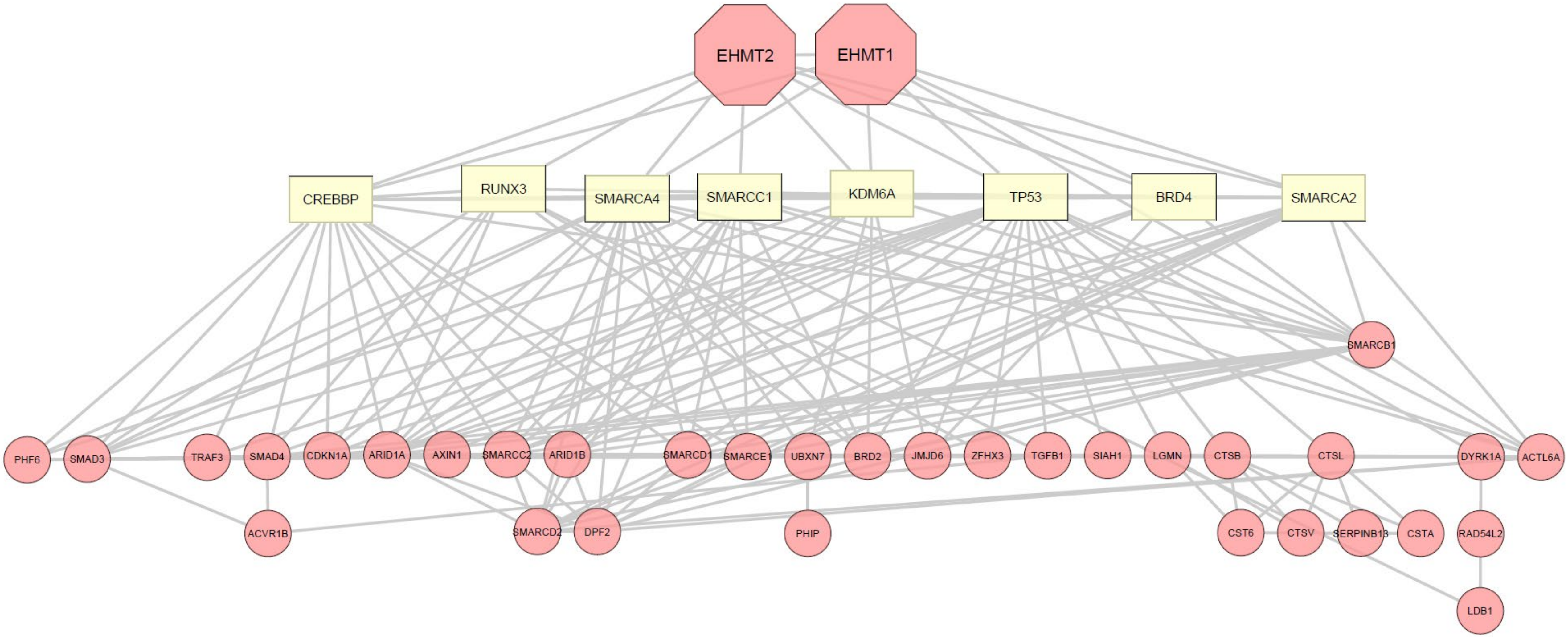
